# Ultra-Fast Multi-Organ Proteomics Unveils Tissue-Specific Mechanisms of Drug Efficacy and Toxicity

**DOI:** 10.1101/2024.09.25.615060

**Authors:** Yun Xiong, Lin Tan, Wai-kin Chan, Eric S. Yin, Sri Ramya Donepudi, Jibin Ding, Bo Wei, Bao Tran, Sara Martinez, Iqbal Mahmud, Hamish I. Stewart, Daniel J. Hermanson, John N. Weinstein, Philip L. Lorenzi

**Author notes:** These authors contributed equally to this work (Yun Xiong, Lin Tan, Wai-kin Chan).

## Abstract

Rapid and comprehensive analysis of complex proteomes across large sample sets is vital for unlocking the potential of systems biology. We present UFP-MS, an ultra-fast mass spectrometry (MS) proteomics method that integrates narrow-window data-independent acquisition (nDIA) with short-gradient micro-flow chromatography, enabling profiling of >240 samples per day. This optimized MS approach identifies 6,201 and 7,466 human proteins with 1- and 2-min gradients, respectively. Our streamlined sample preparation workflow features high-throughput homogenization, adaptive focused acoustics (AFA)-assisted proteolysis, and Evotip-accelerated desalting, allowing for the processing of up to 96 tissue samples in 5 h. As a practical application, we analyzed 507 samples from 13 mouse tissues treated with the enzyme-drug L-asparaginase (ASNase) or its glutaminase-free Q59L mutant, generating a quantitative profile of 11,472 proteins following drug treatment. The MS results confirmed the impact of ASNase on amino acid metabolism in solid tissues. Further analysis revealed broad suppression of anticoagulants and cholesterol metabolism and uncovered numerous tissue-specific dysregulated pathways. In summary, the UFP-MS method greatly accelerates the generation of biological insights and clinically actionable hypotheses into tissue-specific vulnerabilities targeted by ASNase.

## Introduction

The past decade witnessed remarkable technological and scientific strides that paved the way for advances in systems biology(*1, 2*). Innovations in genome sequencing have enabled the generation of large molecular datasets spanning a range of biological systems and matrices, from cells to tissues to biological fluids(*3, 4*). Concurrently, mass spectrometry (MS)-based proteomics has emerged as an essential tool for high-throughput protein-level measurements, becoming indispensable for functional genomics and systems biology(*5–7*).

MS-based proteomics studies can typically be divided into three major steps: sample preparation, MS data acquisition, and data analysis. Whereas data analysis tools have improved significantly in recent years(*8–12*), sample pretreatment and MS-based data acquisition remain limiting steps in proteomics investigation. A variety of sample preparation workflows have been developed for proteomic studies with specific sample types and experimental goals(*13, 14*). However, most workflows are limited in the number of samples that can be processed simultaneously, precluding their application to larger systems biology studies. For MS data acquisition, proteome coverage is a critical challenge due to the wide dynamic range of protein expression levels in cells, ranging from tens to over a million copies per cell. The most common approach to address that complexity has been multi-shot proteomics with offline peptide fractionation(*15–17*), but this approach necessitates extensive instrument time and large sample volumes. Prompted by those limitations, ongoing effort has been allocated to the development of improved single-shot strategies. Recently, narrow-window DIA (nDIA) was reported to achieve the identification of ∼10,000 proteins within 1 h, and ∼7,000 proteins in a 5- min gradient on the Orbitrap Astral MS(*18*).

Here, building on recent advances with nDIA, we introduce a workflow that combines an optimized nDIA-MS method with high-throughput sample preparation to facilitate rapid and in-depth proteome profiling. The method employs 1-Th quadrupole isolation windows and achieves identification of 6,201 and 7,466 proteins with 1- and 2-min LC gradients, respectively. Our streamlined sample preparation incorporates high-throughput tissue homogenization, adaptive focused acoustics (AFA)-assisted proteolysis, and Evotip-accelerated desalting, enabling the processing of up to 96 tissue samples within 5 h. To demonstrate the utility of our approach, we conducted a systems proteomics analysis of 507 biological samples across 13 tissue types to examine the systems pharmacology of L-Asparaginase (ASNase). The results provided a quantitative landscape of 11,472 proteins, uncovering distinct tissue-specific responses that shed new light on mechanisms of ASNase sensitivity, resistance, and toxicity.

## Results

### Identification of 6,201 proteins per minute with UFP-MS

The Orbitrap Astral MS system combines a modified Orbitrap Exploris 480 MS with the asymmetric track lossless (Astral) analyzer(*19*) (Fig. 1a). The Astral analyzer supports parallel MS2 scanning and achieves MS acquisition rates of up to 200 Hz, making it well suited for ultra-high-throughput proteomics analysis(*20, 21*).

**Fig. 1.**
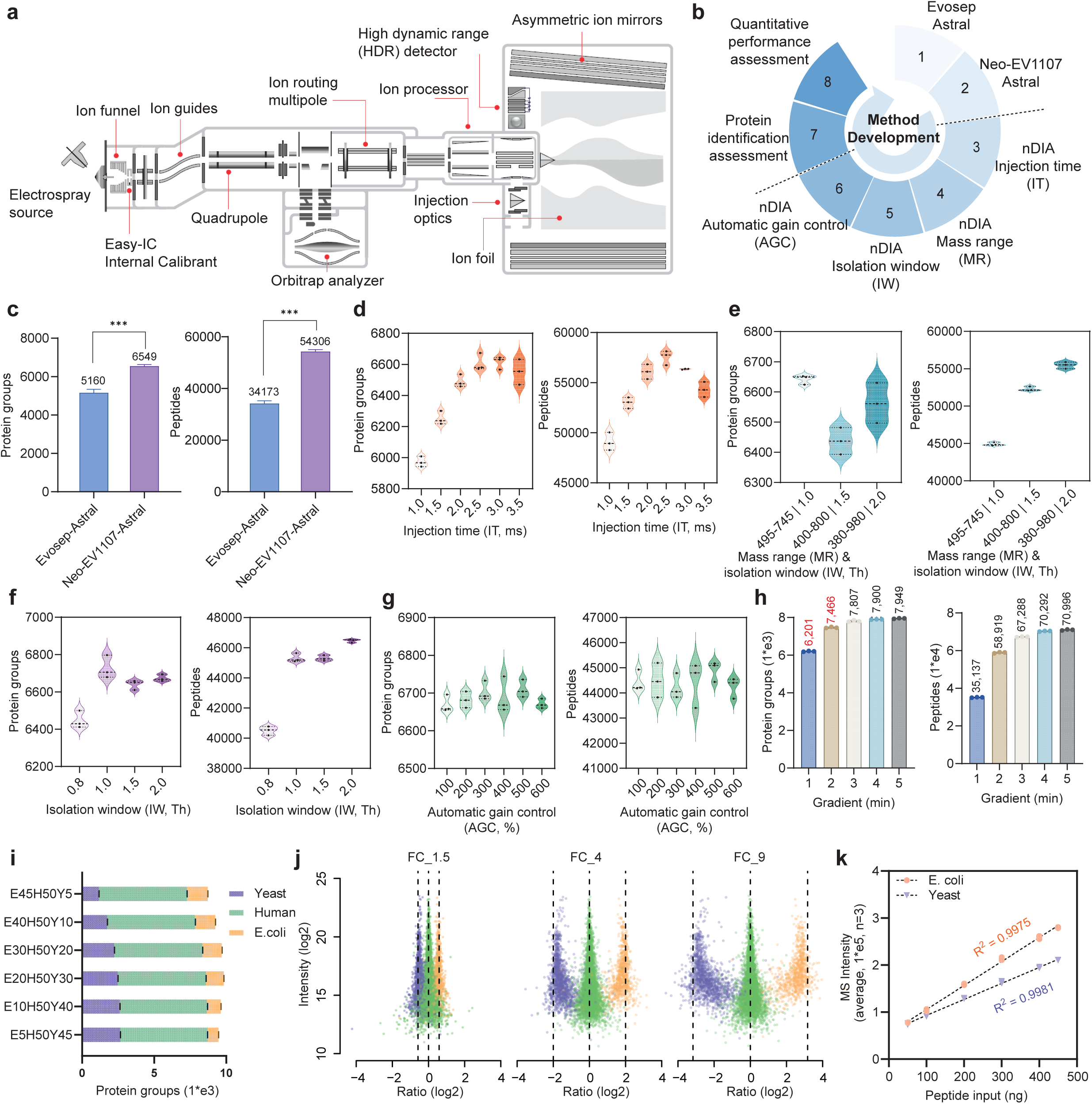
Ultra-fast nDIA enables identification of 6,201 proteins with a one-minute LC gradient **a,** Schematic overview of the Orbitrap Astral MS instrument. **b,** The optimization process of the nDIA method used in this study. **c,** Comparison of the number of identified protein groups and peptides using Evosep-Astral and Neo-Evosep-Astral methods. **d,** Comparison of the number of identified protein groups and peptides with indicated maximum injection time (IT) ranging from 1 ms to 3.5 ms. **e,** Comparison of the number of identified protein groups and peptides with different combinations of mass range (MR) and isolation window (IW). **f,** Comparison of the number of identified protein groups and peptides with MS2 IW ranging from 0.8 Th to 2.0 Th. **g,** Comparison of the number of identified protein groups and peptides with automatic gain control (AGC) value ranging from 100 to 600. **h,** Number of the identified protein groups and peptides using the refined nDIA method with 1- to 5-min LC gradients. **i,** Protein groups identified from mixed samples containing protein digests from *E. coli* (E), human (H), or yeast (Y) in various ratios (E5H50Y45, E10H50Y40, E20H50Y30, E30H50Y20, E40H50Y10 and E45H50Y5). **j,** Peptide Ion intensity as a function of ratio for all runs in (**i**). Dashed lines represent expected log2(A/B) values for proteins from humans (green), yeast (purple) and *E. coli* (orange). FC_9, FC_4, and FC_1.5 are calculated from log2(E45H50Y5/E5H50Y45), log2(E40H50Y10/E10H50Y40), and log2(E30H50Y20/E20H50Y30), respectively. **k,** Correlation between MS intensity and peptide input for *E. coli* and yeast from runs performed in (**i**).

As a benchmark for ultra-high-throughput analysis, the Evosep One system (Evosep, #AN-024A-500SPD) coupled with a timsTOF Pro 2 identifies 2,500 proteins with a 2.2-min LC gradient, achieving a protein identification rate of over 1,000 proteins per min (Supplementary Fig. 1). Based on that, we first coupled an Evosep One system with an Orbitrap Astral MS (Fig. 1b). Using a similar LC gradient and nDIA MS method, we identified an average of 5,160 proteins and 34,173 peptides from 200 ng of HeLa digest (Fig. 1c). To further improve proteome coverage, we modified the LC system by transferring the Evosep endurance column (Evosep, #EV1107, 4 cm x 150 μm, 1.9 μm particle size) and stainless-steel emitter (Evosep, #EV1086, ID 30 μm) to a Vanquish Neo UHPLC system. This modification permitted greater LC gradient flexibility. With an optimized 2-min gradient, proteome coverage was improved to an average of 6,549 proteins and 54,306 peptides (Fig. 1c).

Thereafter, we evaluated various MS data acquisition parameters (Fig. 1b). Consistent with previous studies(*18*), decreasing MS2 maximum injection time (IT) from 3.5 ms to 2.5 ms improved peptide detection (Fig. 1d). Further decreases below 2 ms led to decreased identifications. We next investigated the impact of mass range (MR) on duty cycle, comparing mass ranges of 380-980, 400-800, and 495-745 Th with corresponding isolation windows (Fig. 1e). The 495-745 Th mass range with a 1 Th isolation window delivered optimal results at the protein level. Further exploration of isolation windows from 0.8 to 2.0 Th confirmed that a 1 Th isolation window yielded the highest number of protein identifications (Fig. 1f). Automatic gain control (AGC) target values ranging from 100 to 600 produced similar results, with an AGC of 500 marginally improving precursor ion detection (Fig. 1g). Based on the evaluations, we established a modified nDIA method with an injection time of 2.5 ms, mass range of 495-745, 1 Th isolation window, and an AGC value of 500.

We next assessed the performance across LC gradients from 1 to 5 min. Notably, even with a 1-min gradient, we still identified ∼6,201 human proteins from 1 ug HeLa digest, albeit with a decrease in peptide number (Fig. 1h). A 2-min gradient yielded 7,466 proteins and 59,919 peptides. Proteome coverage remained at nearly 8,000 proteins with gradients from 3 to 5 min. The protein identification rate of up to 6,201 proteins per min greatly exceeds that of previous reports(*22–24*) (Supplementary Fig. 1a, b).

To evaluate the quantitative performance of the new method, we analyzed proteome samples from three species (human, yeast, and *Escherichia coli*) mixed in various ratios. Overall, an average of 9,434 proteins were identified in each mixed sample (Fig. 1i). Quantitative analysis demonstrated accurate protein quantitation across six different ratios (1:9, 1:4, 2:3, 3:2, 4:1, and 9:1) (Fig. 1j). By comparing the different mixtures, we found that the total MS intensities of yeast and *Escherichia coli peptides* increase linearly with respect to sample input (Fig. 1k).

### High-throughput sample preparation accelerates large-scale tissue proteomics

To address the next rate-limiting step, sample preparation, we developed a rapid workflow integrating high-throughput tissue homogenization, adaptive focused acoustic (AFA)-accelerated proteolysis, and Evotip desalting (Fig. 2a). The Precellys Evolution tissue homogenizer performs bead-based protein extraction for up to 24 samples simultaneously, achieving thorough homogenization in less than 1 min. Protein digestion is accelerated using a Covaris R230 Focused-Ultrasonicator, which can handle 96 samples concurrently. Peptide desalting with racked Evotips enabled simultaneous cleaning of up to 96 peptide samples in 30 min.

**Fig. 2.**
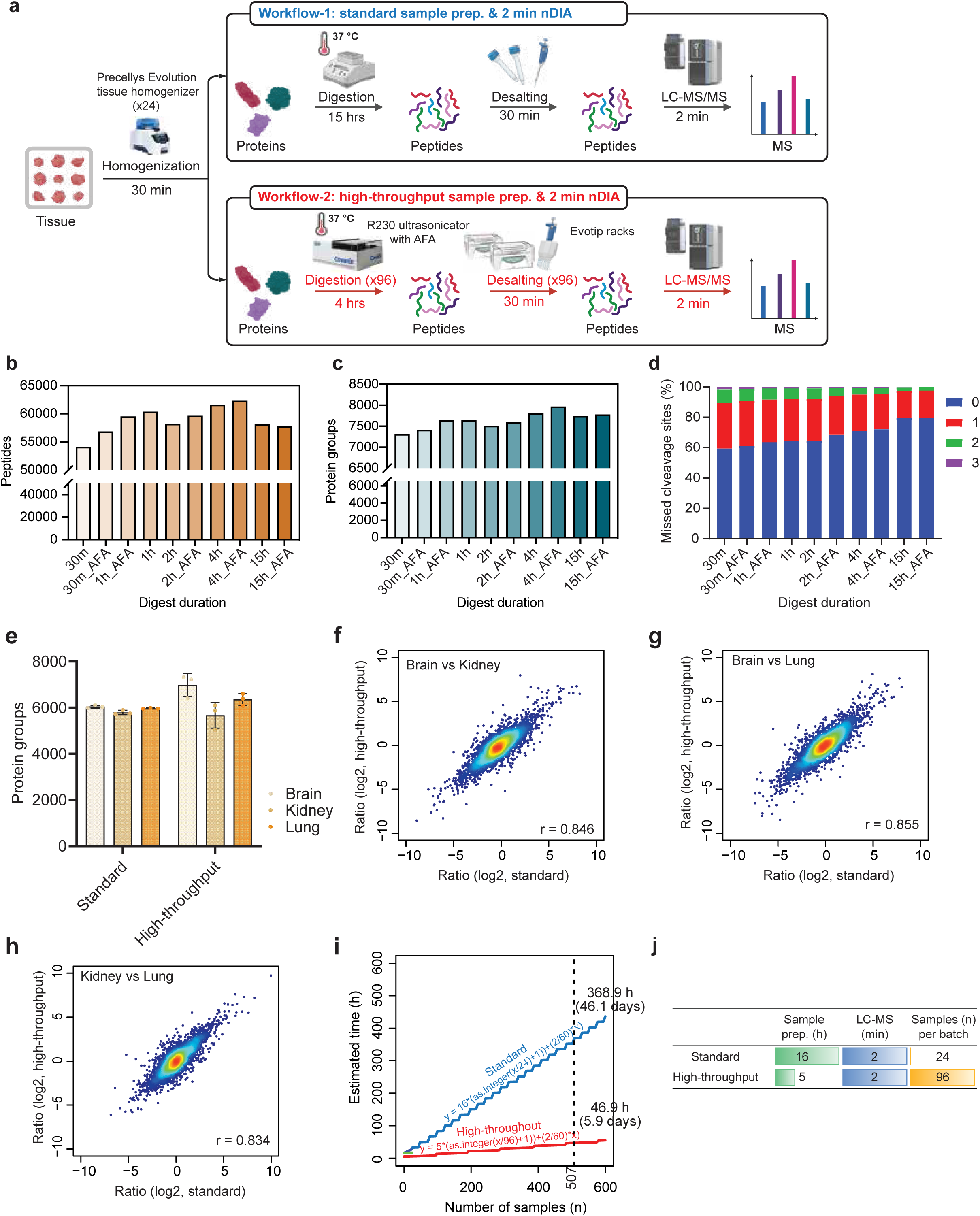
High-throughput sample preparation accelerates large-scale tissue proteomics. **a,** Workflows for standard and high-throughput sample preparation for large-scale tissue proteomics. **b,c,** Number of identified peptides (**b**) and proteins groups (**c**) using indicated proteolytic digestion durations with or without adaptive focused acoustics (AFA)-assisted proteolysis. **d,** Percentages of identified peptides with 0, 1, 2, or 3 missed cleavage sites for indicated proteolytic digestion durations with or without AFA-assisted proteolysis. **e,** Number of identified protein groups with samples prepared from brain, kidney, and lung tissues using the standard and high-throughput workflows. **f-h,** Correlation analysis of the pairwise ratios among three mouse tissues (brain, lung, and kidney) calculated using the data generated by standard and high-throughput workflows. **i,j,** Comparison of the estimated time cost for the standard and high-throughput workflows.

We next compared proteolytic digestion performance with and without sonication across durations from 30 min to 15 h (Fig. 2b-d). MS results showed that 4-h digestion with AFA-assisted sonication provided the highest number of peptides and proteins, although minor differences were observed across conditions (Fig. 2b, c). Over 90% of peptides had only one or no missed cleavage sites across all conditions (Fig. 2d), with AFA-assisted digestion exhibiting slightly better performance in all scenarios. We ultimately chose 4-h digestion to maximize protein recovery, which translates to a total of ∼5 h to process 96 samples (Fig. 2a).

We next analyzed three mouse tissues (brain, kidney, and lung) to compare the output obtained from a standard workflow and the optimized high-throughput workflow (Fig. 2e, Supplementary Fig. 1c). The high-throughput method identified an average of 6,976, 5,665, and 6,355 proteins from brain, kidney, and lung, respectively, which was comparable to the results from the standard workflow. Pairwise comparisons of protein abundance ratios between two tissues at a time demonstrated Pearson correlation coefficients consistently exceeding 0.83 (Fig. 2f-h), indicating comparable quantitative performance between the two methods.

We next estimated the sample preparation time required to process 507 tissue samples (Fig. 2i, j). The standard workflow requires 369 work hours, equivalent to 46.1 x 8-h workdays. The high-throughput workflow, however, needs only 46.9 work hours (5.9 workdays), providing more than seven times higher throughput for large-scale studies.

### Multi-organ quantitative profiling of 11,472 proteins following ASNase treatment

L-Asparaginase (ASNase) is a critical component of the standard of care for pediatric acute lymphoblastic leukemia (ALL). However, its application to adult leukemias and solid tumors is hindered by significant side effects and drug resistance(*25*). ASNase toxicity is attributed in part to its glutaminase co-activity(*25*). There has been interest in engineering glutaminase-free variants that decrease side effects while maintaining efficacy(*26*). We previously demonstrated the anticancer activity of the glutaminase-free Q59L mutant (ASNase^Q59L^) against ASNS-negative cell lines(*27–29*), but those findings failed to translate to a preclinical mouse model(*28*), suggesting that ASNase glutaminase activity is necessary for its anticancer efficacy (Fig. 3a). To aid in the development of new ASNase-based therapeutic strategies, it is essential to gain a systems-level understanding of the mechanisms through which ASNase mediates anticancer activity and toxicity.

**Fig. 3.**
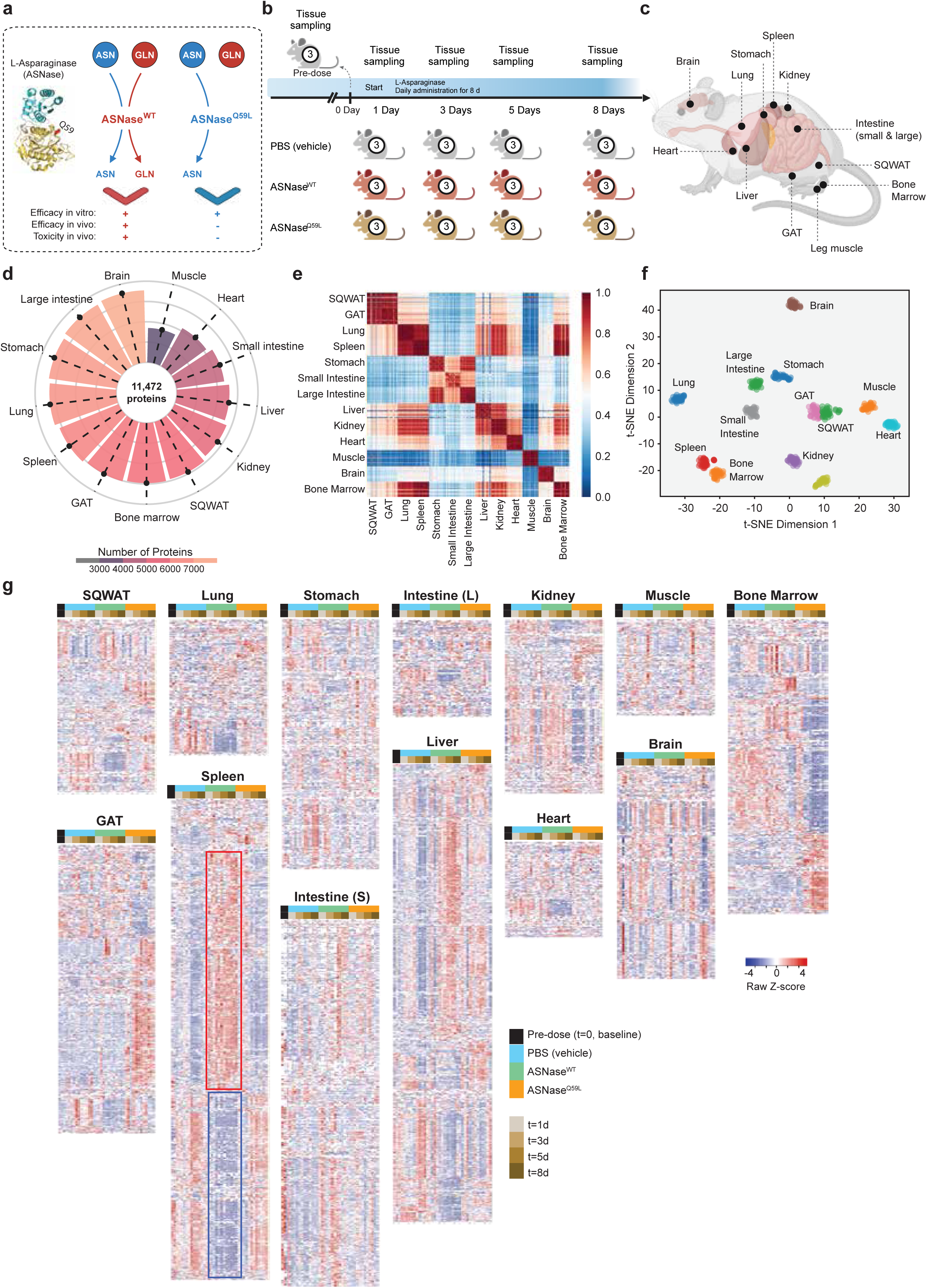
Tissue proteomic landscape of ASNase treatment **a,** Biochemical reactions catalyzed by ASNase^WT^ and ASNase^Q59L^. **b,** Schematic overview of the experimental design. **c,** Overview of the mouse tissues collected for mass spectrometry proteomic analysis. **d,** Total number of proteins identified in each tissue across all treatment groups. **e,** Correlation matrix heatmap of the data generated from 13 mouse tissues. The heatmap was generated using Python. The Pearson correlation coefficient values were calculated based on the MS intensities of each protein in certain samples. **f,** Unsupervised clustering of tissue proteomics data via t-SNE. **g,** Protein abundance in 13 tissues over an 8-d time course of PBS, ASNase^WT^, or ASNase^Q59L^ treatments. t, time.

To that end, we applied the optimized method to analyze tissue samples from NSG mice treated with PBS (vehicle control), 20,000 U/kg ASNase^WT^ (Spectrila®), or 20,000 U/kg ASNase^Q59L^ for 0, 1, 3, 5, and 8 d (Fig. 3b). At each time point, mice were euthanized, and 13 tissues were collected, including subcutaneous white adipose tissue (SQWAT), gonadal adipose tissue (GAT), lung, spleen, stomach, small and large intestine, liver, kidney, heart, leg muscle, brain, and bone marrow (Fig. 3c). Analysis of a total of 507 collected tissue samples yielded 11,472 proteins and 104,784 peptides across all samples (Fig 3d, Supplementary Fig. 1d). The highest numbers of proteins and peptides were observed in the brain, followed by the large intestine, stomach, lung, spleen, GAT, bone marrow, and SQWAT. The lowest identifications were found in leg muscle and heart. As anticipated, the correlation matrix displayed strong associations between the two fat depots, SQWAT and GAT (Fig. 3e). Lung protein expression correlated strongly with that of spleen, bone marrow, and kidney. Large intestine protein expression was more strongly correlated with that of stomach than with small intestine (Fig. 3e). Unsupervised clustering using t-SNE confirmed the similarity of stomach and large intestine (Fig. 3f). We then used heatmaps to illustrate the quantitative overview of protein expression following ASNase treatment in each tissue (Fig. 3g). In most organs, we observed significant differences in protein expression between ASNase^WT^ and PBS treatments, whereas ASNase^Q59L^ yielded only minor differences compared to PBS (Fig. 3g). For example, we saw notable changes in the spleen, but ASNase^Q59L^ also induced unique changes in the bone marrow, kidney, and GAT. These results suggest organ-specific drug effects. We next mined these data to generate new insights into the mechanisms of ASNase efficacy and toxicity across different tissues.

### Amino acid metabolodynamics in whole blood confirm the efficacy of ASNase

To confirm the efficacy of ASNase^WT^, we measured the levels of asparagine (ASN) and aspartic acid (ASP) in whole blood samples (Fig. 4a). Consistent with our previous results(*28, 29*), ASN reached a trough level after one day of treatment with ASNase^WT^ and remained at trough level throughout the 8-day time course, while ASP levels were essentially unaffected (Fig. 4b, c). In contrast, treatment with ASNase^Q59L^ yielded only partial and transient depletion of ASN followed by rapid restoration, suggesting that the body mounted resistance by up-regulating production of ASN.

**Fig. 4.**
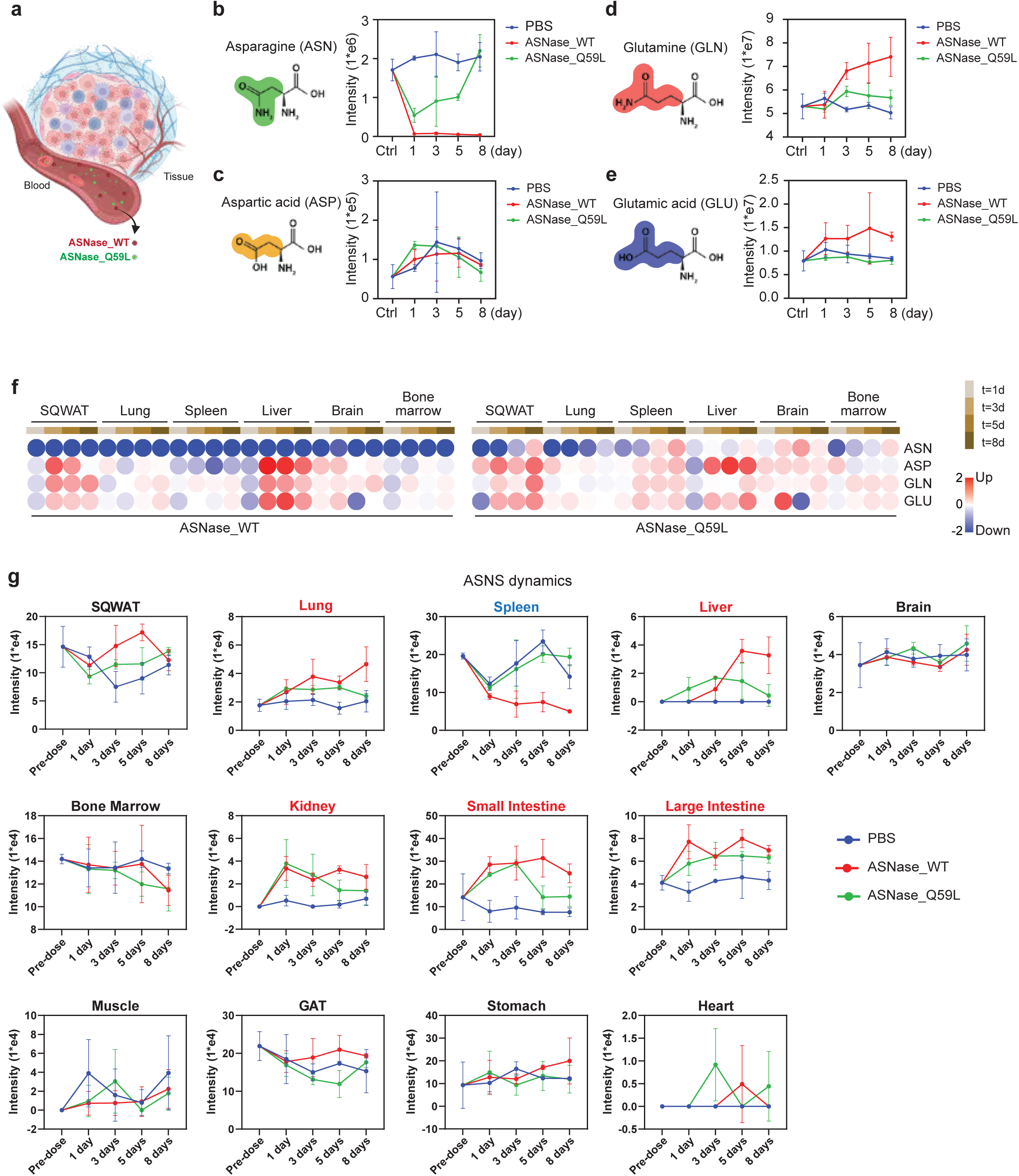
Validation of ASNase efficacy on amino acid metabolism. **a,** System environment architecture of asparaginase (ASNase). **b-e,** The metabolodynamics of asparagine (ASN) (**b**), aspartic acid (ASP) (**c**), glutamine (GLN) (**d**), and glutamic acid (GLU) (**e**) in blood following either PBS, ASNase^WT^, or ASNase^Q59L^ treatment. **f,** The metabolodynamics of ASN, ASP, GLN, and GLU in the six indicated solid tissues following either PBS, ASNase^WT^, or ASNase^Q59L^ treatment. **g,** The dynamic protein levels of ASNS in thirteen indicated solid tissues following either PBS, ASNase^WT^, or ASNase^Q59L^ treatment.

Our previous studies observed that ASNase^WT^ and ASNase^Q59L^ had subtly different effects on glutamine (GLN) and glutamic acid (GLU) levels in a 24-h time course(*28, 29*). In this study, the extended 8-d time course revealed significant increases of both GLN and GLU levels following ASNase^WT^ but not ASNase^Q59L^ treatment (Fig. 4d, e), consistent with the reported up-regulation of glutamine synthetase (GLUL) and glutaminases (GLS, GLS2) following ASNase^WT^ treatment *in vitro*(*30, 31*). These data confirm the *in vivo* efficacy of ASNase^WT^ and highlight key differences between ASNase^WT^ and ASNase^Q59L^.

### Organ-specific resistance and vulnerability to ASNase

To assess the system-wide pharmacodynamic and metabolodynamic effects of ASNase in solid tissues, we monitored ASN, ASP, GLN, and GLU levels across six tissues: SQWAT, lung, spleen, liver, brain, and bone marrow (Fig. 4f). Consistent with observations in blood, ASNase^WT^ treatment resulted in a sustained depletion of ASN in all six tissues, whereas ASNase^Q59L^ initially decreased ASN levels but subsequently up-regulated ASN at these sites (Fig. 4f). We did not observe significant changes in ASP, GLN, or GLU levels after drug treatment in most tissues, except in the liver, where ASNase^WT^ induced increases in all three amino acids after prolonged treatment. Notably, we observed a decrease in ASP levels in the spleen following ASNase^WT^ but not ASNase^Q59L^ treatment. These findings contrast with the observed blood effects, highlighting spleen-specific suppression of ASP metabolism as a targeted vulnerability of ASNase^WT^ treatment.

Asparagine synthetase (ASNS) is a well-established mediator of resistance to ASNase(*25, 27, 28, 32, 33*). However, multiple reports have also described ASNS-low cancer cells exhibiting resistance to ASNase *in vivo*(*28, 34, 35*). In assessing ASNS expression levels we observed rapid and extensive up-regulation in the lung, small intestine, large intestine, liver, and kidney following both ASNase^WT^ and ASNase^Q59L^ treatments (Fig. 4g), suggesting that broad ASNS up-regulation could potentially explain the resistance of low-ASNS tumors to ASNase treatment. In contrast, both ASNase variants suppressed ASNS up-regulation in brain, bone marrow, heart, and muscle. Strikingly, only spleen exhibited down-regulation of ASNS following ASNase^WT^ but not ASNase^Q59L^ treatment. The spleen, a vital organ in the immune system, filters and stores blood and produces white blood cells that protect the body from infection. Given the association of the spleen with blood cells, our findings prompt the hypothesis that spleen-specific suppression of ASNS is a feature of the ASNase mechanism of action against ALL. If future experiments confirm that to be true, combination therapy with an ASNS inhibitor could present a path for extending ASNase treatment to solid tumors.

In addition to ASNS, we examined the effects of ASNase on the broad network of proteins associated with ASN, ASP, GLN, and GLU metabolism (Supplementary Fig. 2a, b). Many of the observed changes suggested adaptive responses to ASNase^WT^. Specifically, we observed up-regulation of: GLUL in spleen, stomach, small intestine, kidney, and heart; GLS in SQWAT, lung, spleen, stomach, small intestine, and kidney; GLUD1 in spleen, stomach, and bone marrow; at least one transaminase (GOT1, GOT2, GPT, GPT2) in all 13 tissues; at least one glutamine transporter (SLC1A5, SLC38A3) in stomach, and liver; and at least one glutathione metabolism component (GCLC, GCLM) in SQWAT, GAT, small intestine, large intestine, brain, and bone marrow. Up-regulation of those glutamine metabolism sub-pathways at the indicated organ sites suggests adaptive resistance mechanisms (i.e., compensatory mechanisms to maintain levels of the four amino acids during nutrient stress).

Nevertheless, five organs exhibited down-regulation of glutamine metabolism in response to ASNase^WT^ treatment (Supplementary Figure 2a, b). SQWAT exhibited down-regulation of CPQ, FOLH1, GOT1, GOT2, and SLC25A22. Spleen exhibited down-regulation of ALDH18A1, CAD, CTPS1, DARS2, GCLM, GMPS, PAICS, PFAS, PPAT, and SLC1A5. Liver exhibited down-regulation of ABAT, CPQ, GGCX, GLS, GLS2, GLUL, GOT2, and NADSYN1. Kidney exhibited down-regulation of ACY1, ACY3, BCAT1, CAD, CTPS1, GATB, GCLC, GPT, and SLC25A22. And muscle exhibited down-regulation of ABAT, ASNS, ASPA, ASS1, BCAT2, CAD, CPQ, DARS2, GCLC, GCLM, GLS, GLUD1, GMPS, GOT1, GOT2, GPT, NARS2, OPLAH, PAICS, PFAS, SLC1A5, and SLC25A22.

Overall, these analyses indicate that most organ sites mount an adaptive response to ASNase^WT^. Nevertheless, potentially vulnerable organs include spleen, SQWAT, liver, kidney, and muscle (Fig. 4g). Among those, spleen uniquely exhibited suppression of ASNS protein (Fig. 4g). No clinical toxicities of ASNase have been associated specifically with spleen, muscle, or kidney, suggesting that using ASNase to treat tumors arising from or supported by those tissues may be safe. Liver and SQWAT were vulnerable organs that have been linked to clinical toxicities.

### System-wide suppression of anticoagulants and cholesterol metabolism as likely drivers of ASNase toxicity

To explore other coordinated stress responses following drug treatment across tissues, we next compared protein expression profiles between ASNase-treated and PBS-treated samples at the pathway level. Proteins with an average fold change in expression greater than 2 or less than 0.5 and a p-value < 0.05 were classified as differentially modulated (Fig. 5a-d, Supplementary Fig. 3-15). ASNase^WT^ treatment significantly modulated (positively or negatively) the vast majority of proteins in liver and spleen (Fig. 5a, b). ASNase^Q59L^ treatment mainly up-regulated proteins in the liver, GAT, and small intestine, while down-regulating proteins in kidney, bone marrow, stomach, liver, and brain (Fig. 5c, d).

**Fig. 5.**
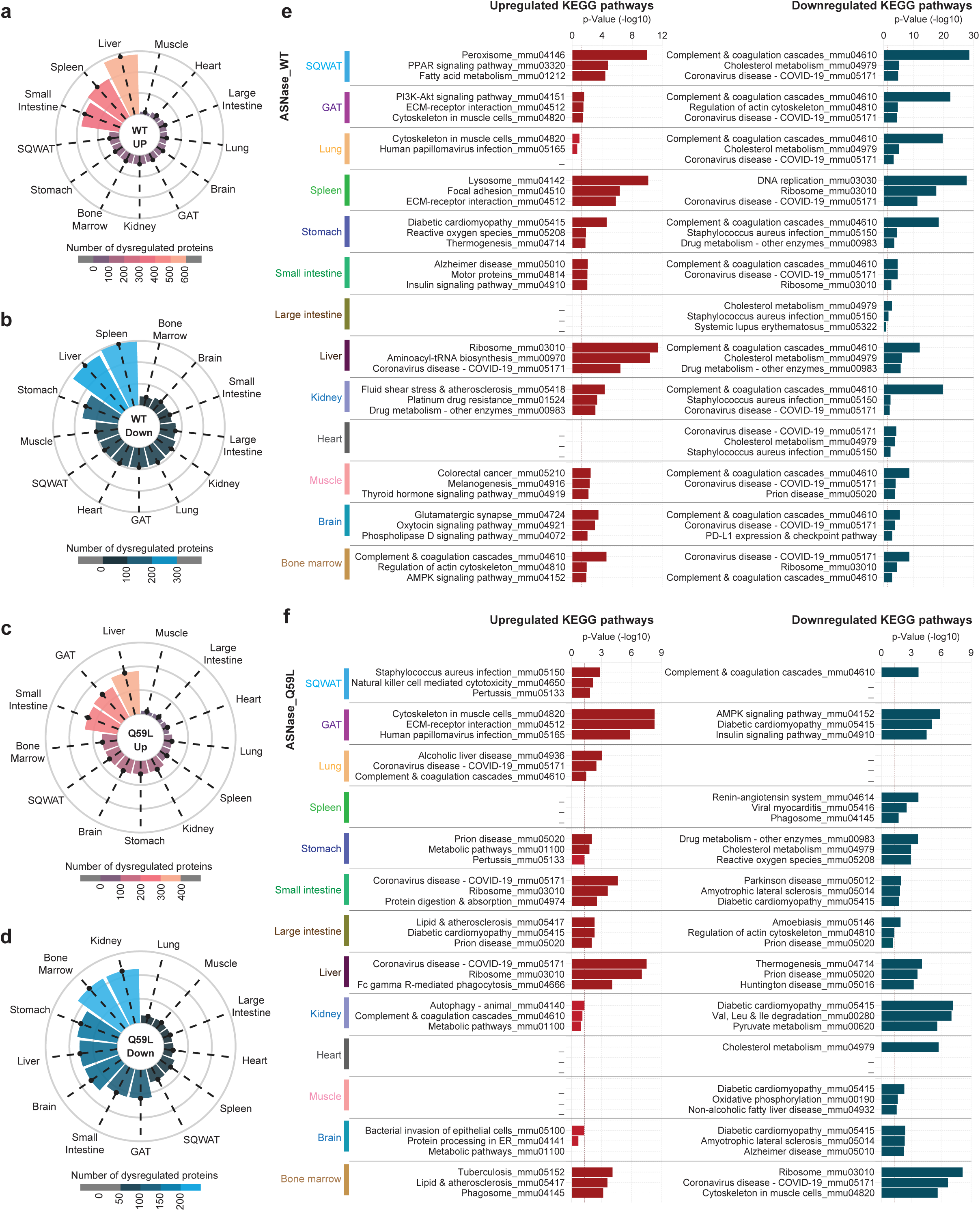
Statistical overview and KEGG pathway enrichment analysis of proteins modulated by ASNase treatment. **a,** Protein expression levels following ASNase^WT^ treatment were compared with PBS-treated controls. Proteins with a fold change greater than 2 and a p-value less than 0.05 were classified as up-regulated. Data from all the time points (1d, 3d, 5d, and 8d) were grouped together. **b,** Same as (**a**) except proteins with a fold change less than 0.5 and a p-value less than 0.05 were classified as down-regulated. **c,** Same as (**a**) except ASNase^Q59L^ compared to PBS treatment. **d,** Same as (**b**) except ASNase^Q59L^ compared to PBS treatment. **e,** KEGG pathway analysis of the up- or down-regulated proteins following ASNase^WT^ treatment in the indicated tissues. **f,** Same as (**e**) except ASNase^Q59L^ treatment.

KEGG pathway enrichment analysis provided several notable insights (Supplementary Fig. 8-12). First, the suppressive effects of ASNase^WT^ were highly consistent across organ sites, with pathways such as “complement and coagulation cascades,” cholesterol metabolism, and coronavirus disease being nearly uniformly downregulated in all tissues (Fig. 5e). No pathways were uniformly up-regulated across all tissues (Fig. 5e), indicating that the up-regulated pathways are highly tissue-specific. Similarly, ASNase^Q59L^ also did not uniformly modulate any pathways across organ sites (Fig. 5f).

Closer inspection of the MS proteomic data confirmed that most identified proteins in the complement and coagulation cascade and cholesterol metabolism pathways were down-regulated across all 13 tissue types following ASNase^WT^ but not ASNase^Q59L^ treatment (Fig. 6a). Focusing on proteins that were consistently dysregulated across multiple tissues, 28 proteins were commonly up-regulated, and 68 proteins were commonly down-regulated across at least three tissues following ASNase^WT^ treatment (Fig. 6b, c). ASNase^Q59L^ treatment up- or down-regulated 56 and 24 shared proteins, respectively (Fig. 6d, e). Around 50% of all proteins commonly down-regulated by ASNase^WT^ were involved in cholesterol metabolism or the complement and coagulation cascades (Fig. 6f-h). Together, these results suggested that well-known ASNase toxicities—hypertriglyceridemia and venous thromboembolism (VTE)—result from system-wide down-regulation of cholesterol metabolism and the complement and coagulation pathways.

**Fig. 6.**
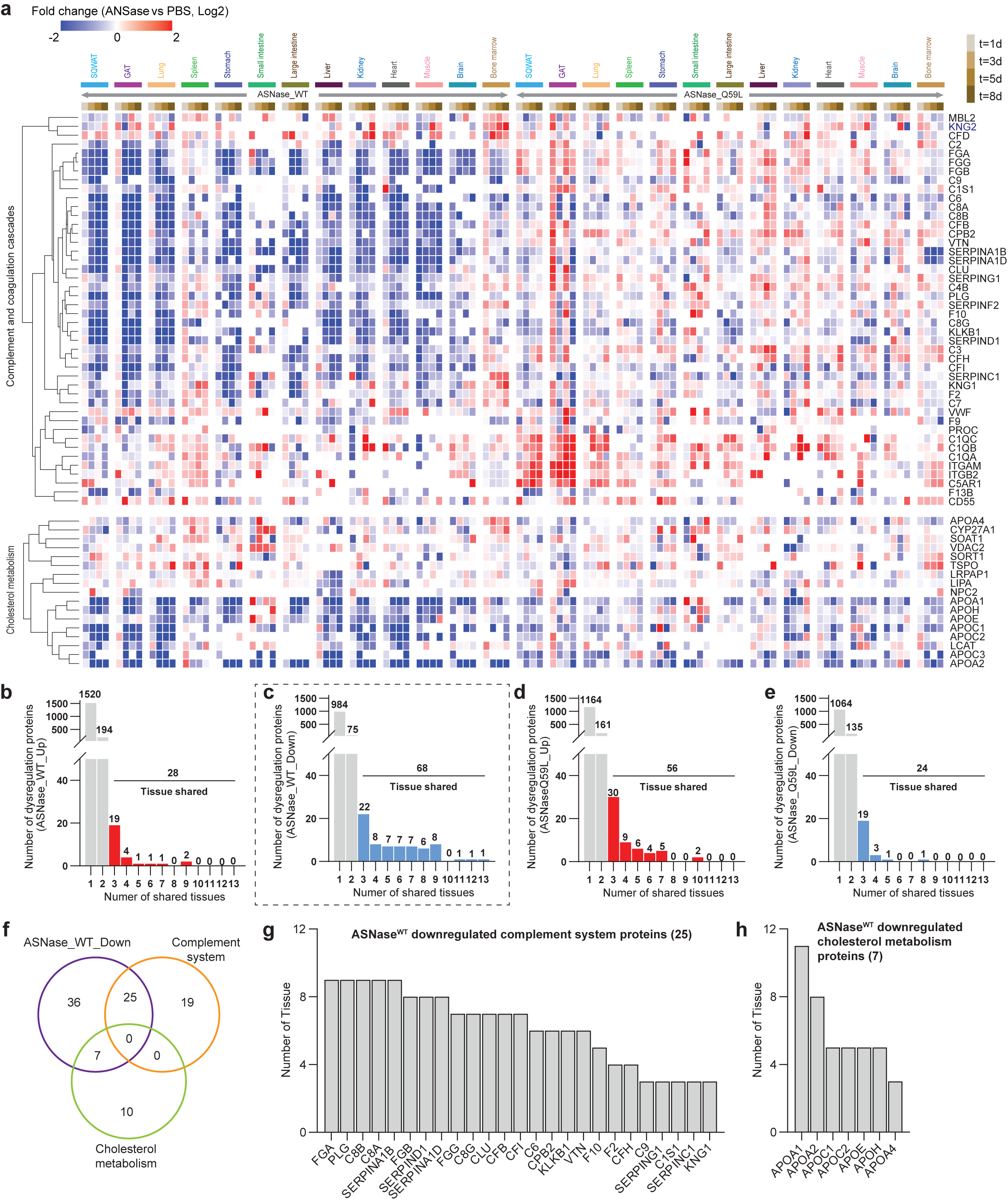
ASNase dysregulates a common set of proteins across multiple tissue types. **a,** Heatmap overview of the dynamic protein levels change of proteins involved in the complement and coagulation cascades (top) and the cholesterol metabolism pathway (bottom) following ASNase^WT^ or ASNase^Q59L^ treatment across all the tested tissues. Protein expression levels following ASNase^WT^ or ASNase^Q59L^ treatment were compared to PBS-treated controls, with the Log2 fold change values presented. **b-e,** Statistical summary of the dysregulated proteins that were common in certain number of tissues. Dysregulation proteins were filtered with a p-value < 0.05 and fold change > 2 or < 0.5. Proteins commonly dysregulated in at least three tissues were colored in red (up-regulation) or blue (down-regulation). **f,** Venn diagram of proteins downregulated by ASNase^WT^, proteins involved in the complement system, and proteins involved in cholesterol metabolism. **g,h,** Downregulated proteins involved in (**g**) complement system or (**h**) cholesterol metabolism following ASNase^WT^ treatment. Only proteins significantly modulated in at least three tissues are displayed.

### ASNase^WT^ modulation of metabolites and proteins converge on inhibition of DNA replication and activation of ferroptosis in spleen

Although down-regulation of cholesterol metabolism and the complement and coagulation pathways was observed in nearly all tissues tested, this was not the case in the spleen (Fig. 5e). As noted previously, spleen uniquely exhibited decreased ASP levels (Fig. 4f) and decreased ASNS protein levels following ASNase^WT^ treatment (Fig. 4g). Considering the strong association between spleen and blood cells, we next analyzed the quantitative MS proteome profiles of the spleen following drug treatment. We compared PBS- and ASNase^WT^-treated spleen proteomes and subjected differentially modulated proteins with a p-value < 0.05 and fold change > 2 or < 0.5 to functional clustering against the KEGG pathways database (Fig. 6a-c)(*36*). Since no dysregulated pathways were enriched prior to 3 d of treatment, we focused on effects at 3, 5, and 8 d. Notably, proteins associated with the “lysosome” pathway were significantly up-regulated at all time points, and DNA replication was significantly down-regulated at all time points (Fig.7a-c). At 8 d, “glutathione metabolism” and “ferroptosis” were significantly up-regulated (Fig. 7c). ASNase^Q59L^ did not affect those pathways (Supplementary Fig. 5).

**Fig. 7.**
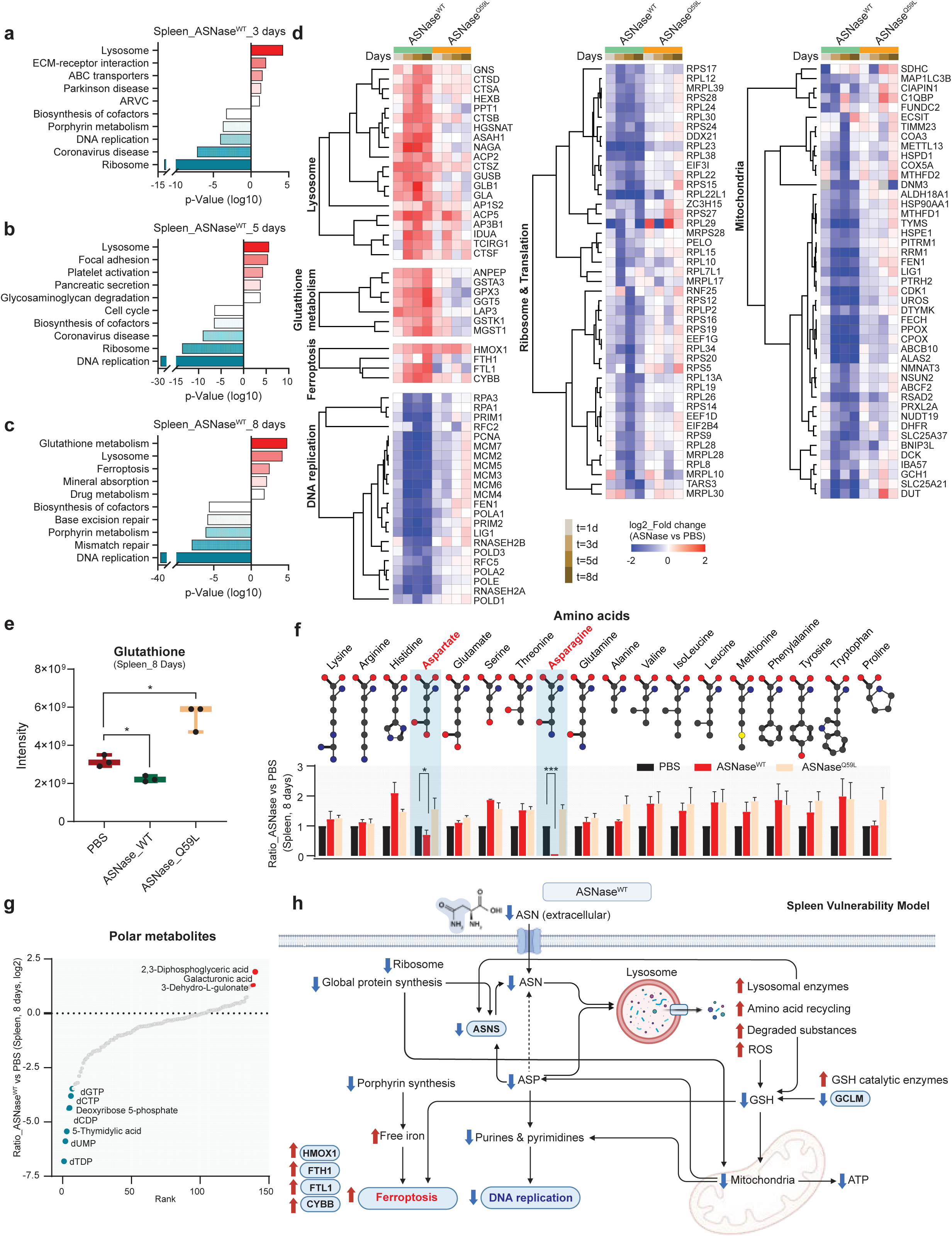
Modulation of the spleen proteome by ASNase. **a-c,** KEGG pathway analysis of the upregulated or downregulated proteins following 3 days (**a**), 5 days (**b**), and 8 days (**c**) ASNase^WT^ treatment in the spleen. **d**, Heatmap overview of the dynamic protein levels change of proteins involved in lysosome, glutathione metabolism, ferroptosis, DNA replication, ribosome and translation, and mitochondria following ASNase^WT^ or ASNase^Q59L^ treatment across all the tested tissues. Protein expression levels following ASNase^WT^ or ASNase^Q59L^ treatment were compared to PBS-treated controls, with the Log2 fold change values presented. **e,** Glutathione levels in spleen samples from mice treated with PBS, ASNase^WT^, or ASNase^Q59L^ for 8 d. **f,** The dynamic regulation of indicated amino acid levels following 8 days treatment with ASNase^WT^ or ASNase^Q59L^ treatment in the spleen. **g,** Polar metabolite levels in spleen samples from mice treated with ASNase^WT^ for 8d. **h,** Model of spleen vulnerability to ASNase^WT^ treatment.

Consistent with previous reports of ASNase-induced autophagy (a lysosome-driven process)(*37*), our data showed up-regulation of: i) lysosomal cysteine proteases CTSA, CTSB, CTSD, CTSF, and CTSZ; ii) lysosomal glycosidases GLA, GLB1, GNS, GUSB, HEXB, and NAGA; and iii) lysosomal phosphatases ACP2 and ACP5 following treatment of ASNase^WT^ (Fig. 7d). ASNase^Q59L^ did not significantly modulate lysosomal proteins. The induction of lysosomal proteins may be a consequence of decreased ASP and ASN, since lysosome-driven autophagy plays an important role in producing amino acids via recycling of proteins(*38*).

Prolonged ASNase^WT^ treatment also up-regulated glutathione (GSH) metabolism and ferroptosis (Fig.7c). In parallel, our metabolomic data revealed that GSH was decreased by ASNase^WT^ and significantly increased by ASNase^Q59L^ in the spleen (Fig. 7e). Most up-regulated GSH metabolism proteins were catalytic enzymes (Fig. 7d), including glutathione peroxidase 3 (GPX3), glutathione hydrolase 5 heavy chain (GGT5), and glutathione transferases (GSTA3, GSTK1, and MGST1). Their up-regulation following ASNase^WT^ treatment was consistent with the decrease of GSH level (Fig. 7e). Since GSH is a key reducing equivalent and negative modulator of ferroptosis, its concurrent down-regulation with the up-regulation of ferroptosis was mechanistically consistent(*39*). The up-regulated ferroptosis pathway encompassed four proteins including CYBB, FTH1, FTL1, and HMOX1 (Fig. 7d). We also observed down-regulation of porphyrin metabolism following ASNase^WT^ for 3 and 8 days (Fig. 7a, c), prompting the hypothesis that down-regulation of porphyrin metabolism permits accumulation of free iron, which is required for ferroptosis (Fig.7c).

A number of DNA replication proteins were down-regulated by ASNase^WT^ treatment (Fig. 7a-d): a) DNA polymerases POLA1, POLA2, POLD1, POLD3, and POLE; b) components of the MCM2-7 complex MCM2, MCM3, MCM4, MCM5, MCM6, and MCM7; c) DNA primases PRIM1 and PRIM2; d) ribonuclease H2 subunits RNASEH2A and RNASEH2B; and e) other DNA replication-associated proteins PCNA, RPA1/3, RFC2/5, FEN1, and LIG1 (Fig. 7d). Down-regulation of DNA replication may be associated with the decrease of ASP (Fig. 4f), which contributes to *de novo* synthesis of purine and pyrimidine nucleotides(*40*). Interestingly, of all amino acids in spleen following ASNase treatment, only ASP and ASN were clearly down-regulated (Fig. 7f, Supplementary Fig. 16). Correspondingly, metabolomics indicated significant decrease of several deoxyribonucleotides, including dTDP, dUMP, dTMP, dCDP, dCTP, and dGTP (Fig.7g), further supporting the observed down-regulation of nucleotide synthesis at the protein and pathway levels.

Spleen-specific down-regulation of ASNS may be associated with up-regulation of aminopeptidases LAP3 and ANPEP (Fig. 7d) and down-regulation of ribosome and translation pathways (Fig. 7a, b, d, Supplementary Fig 17a, b). A number of mitochondrial proteins that were specifically down-regulated in spleen (Fig. 7d, Supplementary Fig.17a, b), which is consistent with previous reports that mitochondrial electron transport chain enables synthesis of ASP(*41*). To note, in addition to the deoxyribonucleotides that described above, adenosine triphosphate (ATP) was also decreased in spleen following ASNase^WT^ treatment (Supplementary Fig 17c).

Collectively, integrated analysis of proteomic and metabolomic data implicates the spleen as a highly vulnerable target of ASNase^WT^. That vulnerability is potentially explained by signaling cascades that modulate amino acid metabolism, energy metabolism, redox metabolism, and global protein synthesis converging on inhibition of DNA replication and induction of cell death (ferroptosis) (Fig. 7h), suggesting a plausible mechanism supported by our findings: 1) ASNase^WT^ decreases ASN and ASP in spleen (Fig. 7f); 2) endoplasmic reticulum stress and protein misfolding are up-regulated(*42, 43*)); 3) lysosomal enzymes are activated (Fig. 7d); 4) reactive oxygen species (ROS) accumulate; 5) GSH is consumed (Fig. 7d, e); 6) damaged mitochondria accumulate; 7) mitophagy clears damaged mitochondria (Fig. 7d) and further decreases ASP production and nucleotide synthesis (Fig. 7f, g) and ATP levels (Supplementary Fig 17c); 8) DNA replication is suppressed (Fig. 7a-d); 9) recycling of damaged RBCs together with suppression of porphyrin synthesis leads to accumulation of free iron (Fig. 7a, c); and 10) ferroptosis is activated (Fig. 7c, d). The observed decrease of ASNS expression is likely due to the combined impairment of protein synthesis (Fig. 7d), decrease of ASP (substrate of ASNS) availability (Fig. 7f), and increased lysosomal degradation (Fig. 7d).

### ASNase dysregulates system-wide and tissue-specific pathways

In addition to findings in spleen, our analysis revealed numerous tissue-specific dysregulated pathways in other tissues following ASNase treatment. These observations offer further insights into the effects of ASNase^WT^ at the system-wide and tissue-specific levels. Those results are discussed in the Supplementary Information.

## Discussion

MS-based proteomics has become an essential tool for high-throughput investigation of cellular proteins. While RNA sequencing technologies are adept at quantifying gene expression levels, the direct analysis of proteins—the principal executors of cellular functions—remains indispensable in biological research.

To improve upon the latest advances of ultra-high-throughput LC (Evosep, #AN-024A-500SPD) and MS (nDIA on Orbitrap Astral MS(*44*)) technologies, we present a novel LC-MS approach that integrates optimized nDIA with short-gradient micro-flow chromatography, yielding identification of 6,201 and 7,466 proteins with 1- or 2-min LC gradients, respectively. The achieved protein identification rate of up to 6,201 proteins per min represents more than a two-fold improvement over previously reported cutting-edge methods (Supplementary Fig. 1a, b)(*18, 22–24, 45*).

Sample preparation also represents a critical rate-limiting step for large-scale proteomics experiments. To address this, we designed a rapid tissue proteomics workflow capable of processing up to 96 tissue samples within 5 h, including a 4 h proteolysis step. We used the workflow to obtain more than seven-fold increased throughput (Fig. 2i) in a multi-organ systems proteomics project involving 507 tissue samples (Fig. 2i). Decreasing the protein digestion time to 30 min increases overall processing speed to 96 samples per hour but results in a marginal decrease (2.2%) in protein identification (Supplementary Table 8). Our chosen application was tissue proteomics, but the workflow can be adapted to other biological matrices.

We chose the systems pharmacology of ASNase as an application primarily because its toxicity has prevented application to adult patients and cancers beyond pediatric ALL(*25*). Additionally, despite >50 years of research, the effects of its glutaminase activity and the mechanistic underpinnings of ASNase sensitivity and resistance are poorly understood.

The quantitative profile of 11,472 proteins provides a detailed multi-organ roadmap of dysregulated cellular pathways following ASNase treatment. New insights regarding ASNase toxicities included the association of VTE with suppression of the complement and coagulation system, and association of hypertriglyceridemia with widespread suppression of cholesterol metabolism.

The complement and coagulation pathways are crucial for host defense against pathogen infection and injury, involving blood-based defense systems that are proteolytically activated to produce active molecules crucial for immunity and clotting(*46*). Genes associated with the coronavirus disease pathway also include complement factors and immune response elements. Within the 44 down-regulated proteins associated with the complement and coagulation cascades, 11 have anti-coagulant function (CPB2, KNG1, KNG2, PLG, PROC, SERPINA1B, SERPINA1D, SERPINC1, SERPIND1, SERPINF2, SPERING1), and the remaining 33 have pro-coagulant function. Most of the 11 anti-coagulant proteins were down-regulated in up to 9 of the 13 tissue types (Fig. 6a). Similarly, previous research on the VTE side effects of ASNase identified down-regulation of plasminogen (PLG), antithrombin III (SERPINC1), and fibrinogen (FGA, FGB, FGG) (*47, 48*). SERPINC1 is of primary interest in current clinical practice; guidance recommends weekly monitoring of SERPINC1 levels in patients and administration of recombinant SERPINC1 when levels fall below 60% with a repletion target of 80% to 120%(*49*). Our findings reveal that SERPINC1 was widely down-regulated in most of the 13 tissues (Fig. 6a) and exhibited a significant decrease in three specific tissues including stomach, liver, and kidney (Fig. 6h). Additionally, PLG, SERPINA1B (human homolog SERPINA1), SERPINA1D (human homolog SERPINA1), and SERPIND1 were even more significantly down-regulated than SERPINC1 (Fig. 6g), representing potentially more impactful targets for mitigating ASNase-induced VTE. Although no recombinant variants of PLG or SERPIND1 exist yet, INBRX-101, a recombinant Fc fusion protein of SERPINA1 (alpha-1 antitrypsin) is entering Phase 2/3 testing(*50*). Given that only 60% of patients with VTE respond to SERPINC1 repletion, our results provide a rationale for testing INBRX-101 for further mitigation of ASNase-induced VTE. All in all, our data confirm previous findings and identify additional changes in complement and coagulation cascade proteins following ASNase treatment, further linking the down-regulation of these pathways to VTE.

Within the 17 down-regulated proteins in the cholesterol metabolism pathway, 6 (APOA1, APOA2, APOA4, APOC2, LCAT, and SORT1) are negative modulators of triglyceride levels, and their down-regulation potentially explains ASNase-induced hypertriglyceridemia(*51, 52*). It has been hypothesized that ASNase-induced hypertriglyceridemia, which occurs in up to 50% of patients, is the result of very low-density lipoprotein (VLDL) up-regulation(*53, 54*). That hypothesis is not sufficiently granular, though, because VLDL components can be positive or negative modulators of triglyceride levels based on whether they facilitate secretion or uptake of triglycerides. Considering VLDL components as a whole, our data revealed prominent down-regulation of APOA4, APOC1, APOC2, APOE, and APOH in SQWAT, lung, liver, large intestine, and heart tissues (Fig. 5e, Fig. 6a, c, h). All of those except APOE are negative modulators of triglyceride levels; their down-regulation could potentially explain ASNase-induced hypertriglyceridemia. In four tissues we also observed a notable increase in adiponectin (ADIPOQ), which promotes the catabolism of VLDL triglycerides and may explain the observed down-regulation of VLDL apolipoproteins (Fig. 6b)(*55*). Together, these findings support the following mechanistic hypothesis to explain ASNase-induced hypertriglyceridemia: i) nutrient stress triggers lipogenic processes culminating in triglyceride synthesis; ii) ASNase suppresses the capacity of organs (primarily the liver) to assemble triglyceride-VLDL complexes, and/or ASNase-induced ADIPOQ rapidly catabolizes such complexes before they can be exported; and iii) free triglycerides are secreted into systemic circulation without a mechanism to distribute them into tissues (mainly fat), leading to triglyceride accumulation in the blood.

In considering therapeutic strategies to mitigate ASNase-induced hypertriglyceridemia, our results identify ADIPOQ as one potential target, but no inhibitors are currently available. Additional candidate targets include the two known positive modulators of triglyceride levels— APOC3 and APOE. Indeed, our data indicates that APOC3 was maintained near baseline levels or elevated in SQWAT, spleen, stomach, large intestine, kidney, muscle, brain, and bone marrow (Fig. 6a). APOC3 was recently identified as a target for lowering triglycerides by up to 77% using the antisense inhibitor volanesorsen(*56*). APOE, as noted previously, was down-regulated in most tissues, but it was strikingly up-regulated in the small intestine by ASNase. Since APOE-targeted drugs (pioglitazone, rosiglitazone) are available, our findings provide a rationale for testing these and volanesorsen for mitigation of ASNase-induced hypertriglyceridemia.

New insights regarding ASNase efficacy suggested that the spleen and kidney may represent targetable organ vulnerabilities. In response to ASNase^WT^ treatment, spleen exhibited suppression of ASNS, suppressed glutamine metabolism including ASP production, impaired DNA replication, and induction of ferroptosis. Those observations prompt the hypothesis that ASNase may be effective against a broad range of hematological-related malignancies. Although kidney exhibited up-regulation of ASNS, it remains a potential target due to broad suppression of glutamine metabolism induced by ASNase^WT^ treatment. Future work will be required to test the hypothesis that ASNase is effective for treating renal cancers.

New insights regarding ASNase resistance identified system-wide up-regulation of ASNS as a key resistance factor, building upon the reported association between the tumor microenvironment and ASNase resistance(*25, 57*). The observed up-regulation of liver protein synthesis may also contribute to ASNase resistance. Importantly, our findings explain how low-ASNS tumors can still be resistant to ASNase; if ASNase dosage is insufficient to maintain durable depletion of ASN, tumor proliferation can continue. Because virtually every organ throughout the body up-regulates ASNS to produce ASN following ASNase treatment, the battle between the drug and the body can only be won if ASN depletion is sustained. These findings highlight the critical need for new therapeutic drug monitoring protocols focused on ASN measurement in whole blood, rather than relying on methods that measure only enzyme activity levels.

The use of ASNS as a predictive biomarker of response to ASNase therapy presents complex considerations. Our findings indicate that ASNase suppresses ASNS up-regulation in selected tissue types, and tumors that exhibit that phenotype are likely to respond well to ASNase. However, because most tissue types exhibit significant up-regulation of ASNS in response to ASNase treatment, tumor-specific ASNS expression may play a relatively minor role in ASNase resistance. Until further clarity is achieved, efforts to extend ASNase therapy to new indications should continue to prioritize testing against tumors with low ASNS expression.

Finally, we hasten to emphasize that integrated analysis of amino acids—particularly ASN, ASP, GLU, and GLN—and their associated proteins proved to be particularly valuable for characterizing organ-level nutrient dependencies.(*58*)

In summary, ultra-fast, quantitative proteome profiling enabled deep analysis of the systems pharmacology of ASNase, facilitating the generation of novel insights into the enzyme-drug’s efficacy and toxicity. This work highlights the transformative potential of nDIA-based ultra-fast proteomics and demonstrates its applicability to a broad range of life science investigations, including the systems biology of dietary interventions, exercise interventions, environmental factors, genetics, and beyond.

## Acknowledgments

We thank all members of MD Anderson Proteomics Core Facility and Metabolomics Core Facility for their help and constructive discussions.

## Author contributions

Y.X., W.K.C., and P.L.L. conceived the project. Y.X. performed all experiments with assistance from L.T., W.K.C., E.S.Y., S.R.D., J.D., B.W., B.T., S.M., I.M., H.I.S., and D.J.H. L.T., B.T., and S.M. conducted the metabolomics analysis. W.K.C., E.S.Y., and S.R.D. carried out the mouse experiments. Y.X. handled data analysis and figure preparation. Y.X. wrote the manuscript, while P.L.L. and J.N.W. revised it with input from all authors. P.L.L. and J.N.W. supervised the study.

## Data and materials availability

All data are available in the main text or the supplementary materials. The mass spectrometry proteomics data have been deposited to the ProteomeXchange Consortium via the PRIDE partner repository with the dataset identifier PXD055919 and PXD055590.

## Supplementary Materials

Supplementary Results

Supplementary Discussion

Online methods

Supplementary Figs. 1 to 19

Supplementary Tables 1 to 27

References (1-35)

## Supplementary Materials

### Supplementary Results

#### ASNase dysregulates system-wide and tissue-specific pathways

In addition to its broad impact on lipid metabolism and the complement system, ASNase^WT^ induces specific dysregulation of various pathways in different tissues. Notably, while ribosomal proteins were reduced in the spleen, small intestine, and bone marrow, ASNase^WT^ treatment led to a significant increase in proteins associated with ’ribosome,’ ’ribosome biogenesis,’ and ’aminoacyl-tRNA synthesis’ in the liver (Supplementary Fig. 18a). In parallel, drug-treated liver exhibited significant up-regulation of amino acids (Supplementary Fig. 16b). This suggests heightened protein biosynthesis activity in the liver. Furthermore, purine metabolism and RNA polymerases were also upregulated by ASNase^WT^ at 8 d (Supplementary Fig. 18a), indicating elevated gene expression in the drug-treated liver. These observations are consistent with the well-known function of liver in synthesizing plasma proteins(*1*), and it is not surprising to see evidence of the liver remaining a protein synthesis factory in the presence of significant nutrient stress.

Bone marrow serves as the microenvironment for leukemias and is known to protect leukemia cells against ASNase by secreting ASN to feed the cancer cells or lysosomal cysteine proteases to degrade and clear ASNase(*2*). We observed significant bone marrow up-regulation of cysteine proteases including CTSC, CTSG, and CTSL after 8 d of ASNase^WT^ treatment (Supplementary Fig. 18b). ASNase^Q59L^ also induced up-regulation of CTSC and CTSG (Supplementary Table 11). KEGG pathway enrichment analysis revealed up-regulation of the “complement and coagulation cascades” pathway in the bone marrow—the opposite direction observed in other tissues. Additionally, proteins involved in “AMPK signaling” and “PPAR signaling” were also induced, including ADIPOQ, SCD1, PCK1, and PIK3R2. These observations are consistent with previous reports that ASNase treatment or ASNS knockdown activated AMPK(*3*), which is a negative modulator of TORC1 and a positive modulator of autophagy, which connects to the observed up-regulation of lysosomal proteases.

ASNase^WT^ treatment induced distinct responses in the two adipose tissue depots, The gonadal adipose tissue (GAT) exhibited induction of “PI3K-Akt signaling,” “ECM-receptor interaction,” and “Focal adhesion” pathways (Supplementary Fig. 18d). The proteins in those pathways are largely overlapping, including COL4A6, COL6A5, TEK, THBS1, and MCL1. TEK, THBS1, and COL4A6 have also been categorized under “PI3K-Akt-mTOR signaling” according to WikiPathways(*4*). The subcutaneous white adipose tissue (SQWAT) exhibited up-regulation of multiple proteins involved in PPAR signaling, including SLC27A1, SLC27A4, ACAA1A, ACAA1B, ACSL1, EHHADH, and PCK1 (Supplementary Fig. 18c), which are primarily associated with fatty acid metabolism. SQWAT is the largest fat depot in most individuals, a critical energy storage depot, and, consequently, known to be more metabolically active than GAT. Hence, the up-regulation of fatty acid metabolism likely represents an adaptive response to nutrient stress induced by ASNase^WT^.

Notably, the “ECM-receptor interaction” and “Focal adhesion” pathways were up-regulated in several tissues, including GAT, lung, stomach, spleen, and liver (Supplementary Fig. 18e). These pathways involve various collagen subunits (COL1A1, COL1A2, COL4A2, COL4A6, COL6A1, COL6A2, COL6A3, and COL6A5) and integrins (ITGA1, ITGA7, ITGA8, ITGAV, ITGB4). ASNase has been reported to disrupt glycosylation, affecting interactions between microvascular endothelial cells, ovarian cancer cells, and ECM components, thereby inhibiting cancer-induced angiogenesis and cancer invasion of surrounding tissues(*5*). In another study, PEGylated ASNase was associated with liver fibrosis due to excessive extracellular matrix (ECM) accumulation(*6*). In ASNase-treated liver, our metabolomic data revealed an increase in 4-hydroxyproline, a major component of collagen that plays a key role in collagen stability(*7*) (Supplementary Fig. 19). Modulation of ECM components may also be associated with inhibition of metastasis by ASNase treatment and ASNS knockdown(*8*).

Another side effect of ASNase is abdominal or stomach pain. Our proteomic data showed up-regulation of “reactive oxygen species (ROS)” and “oxidation phosphorylation” in stomach (Supplementary Fig. 18f). Proteins associated with these categories include mitochondrial inner membrane proteins, such as NADH dehydrogenase subunits (NDUFB3 and NDUFB6), cytochrome c oxidase subunit 6C (COX6C), ATP synthase subunit f (ATP5MF), voltage-dependent anion-selective channel protein 2 (VDAC2), and GTPase NRAS (NRAS).

Like the situation in spleen, glutathione metabolism-related proteins, including various glutathione S-transferases (GSTA1, GSTA3, GSTO1, and GSTT2), were up-regulated in the kidney (Supplementary Fig. 18g). These GSTs are included in the top-3 KEGG pathways induced by ASNase^WT^ in the kidney (Supplementary Fig. 9). ASNS was increased by both ASNase variants in kidney, alongside an increase of several other proteins (PSAT1, GPT2, and PHGDH) involved in amino acid biosynthesis (Supplementary Fig. 18g).

The mTOR signaling pathway was notably induced in ASNase^WT^-treated small intestine (Supplementary Fig. 18h), with key proteins including GTPase KRas (KRAS), dual specificity mitogen-activated protein kinase kinase 1 (MAP2K1), eukaryotic translation initiation factor 4E (EIF4E), insulin receptor subunit alpha (INSR), V-type protein ATPase catalytic subunit A (ATP6V1A), segment polarity protein disheveled homolog DVL-2, large neutral amino acid transporter small subunit 1 (SLC7A5), and calcium-binding protein 39 (CAB39), which binds and activates STK11/LKB1. We also observed massive up-regulation of amino acid metabolism-related sub-pathways including arginine and proline metabolism, beta-alanine metabolism, and aminoacyl-tRNA biosynthesis, which was also induced in ASNase^WT^-treated liver. ASNase^WT^ also induced N-glycan biosynthesis in the small intestine.

In conjunction with elevated Wnt signaling, pharmacologic inhibition of glycogen synthase kinase-3 alpha (GSK3A) was reported to re-sensitize ASNase-resistant leukemias to ASNase through inhibition of proteasome activity(*9*). Here, we found notable induction of Wnt signaling pathway in ASNase^WT^-treated muscle (Supplementary Fig. 18i); the underlying proteins were glycogen synthase kinase-3 beta (GSK3B), catenin beta-1 (CTNNB1), and cAMP-dependent protein kinase catalytic subunit beta (PRKACB). Because our other analyses identified muscle as a targetable vulnerability of ASNase^WT^, these findings suggest that future efforts to test ASNase against sarcomas should include combination with GSK3A inhibitors.

Despite the inability of ASNase to cross the blood-brain barrier, decreased blood ASN levels can lead to decreased ASN levels in cerebrospinal fluid(*10*). Indeed, ASNase toxicity on the central nervous system is well-documented(*11*). Our data showed up-regulation of several pathways in ASNase^WT^-treated brain, with “glutamatergic synapse” being the top-ranked category (Supplementary Fig. 18j). This pathway involves glutamate receptors and signaling proteins, including metabotropic glutamate receptor 3 (GRM3) and other synaptic signaling proteins. These observations may offer insight into ASNase-induced CNS toxicity.

The lung, large intestine, and heart did not reveal significant pathway modulation by ASNase (Supplementary Fig. 4, 11, 15). However, NOTCH1 and several ECM components (COL6A2, COL6A6, and FBN1) were increased following ASNase^WT^ treatment in lung (Supplementary Table 15).

### Supplementary Discussion

It is worthwhile to briefly review the history of rapid sample preparation and fast MS acquisition methods, as our work builds upon these previously reported advancements. As described, sample preparation and MS data acquisition are becoming the limiting steps of MS-based proteomics studies, due to the rapid development in data analysis tools(*12–16*). Many different workflows have been developed for specific sample types and experimental goals(*17, 18*). One attempt to address the issue is the In-StageTip method, which facilitates high-throughput preparation of eukaryotic cell samples but requires a custom, 96-well StageTip holder(*19*). Pressure cycling technology (PCT) requires ∼3h to process 16 samples in one batch(*20*). Adaptive focused acoustics (AFA) technology for rapid sample processing in FFPE tissue proteomics has shown significant promise(*21*). Another promising approach is a simple and efficient workflow that enables simultaneous preparation of up to 192 cell lysate samples in less than 7 h for proteomics analysis, and this method is also suitable for plasma samples(*22*). Despite these advances, there is still an urgent need for faster sample preparation methods that can keep pace with the latest analytical technologies.

In addition to throughput, proteome coverage is another critical challenge in proteomic research due to the wide range of protein expression levels in cells, ranging from tens to over a million copies per cell. To address this complexity, offline peptide fractionation with strong cation exchange (SCX) chromatography, high-pH C18, or hydrophilic interaction liquid chromatography (HILIC), has traditionally been employed to reduce sample complexity and enhance proteome coverage in MS analyses(*23–25*). Yet, this approach necessitates extensive instrument time and large sample volumes to generate and analyze multiple fractions per sample, limiting its scalability in high-throughput systems biology studies, particularly those with restricted sample availability. Prompted by those limitations, single-shot analyses strategies leveraging improved MS data acquisition speeds and sensitivity are becoming more popular. The BoxCar MS1 profiling method quantifies >10,000 proteins in 100 min using a Q Exactive HF MS with the need of a pre-compiled spectral library(*26*). Another group identified 10,044 proteins in 120 min using DIA-based single-shot proteomics on an Orbitrap Exploris 480 MS coupled with FAIMS(*27*). The Thin-diaPASEF method successfully detected up to 11,698 unique proteins from human cancer cell lines with a 100 min active gradient using timsTOF HT(*28*). Recently, several studies measured ∼10,000 identified proteins within one hour using Orbitrap Astral MS(*29–31*).

Leveraging the high scan speed of TripleTOF 6600 mass spectrometer, scanning SWATH (sSWATH) method was reported to enable ultra-fast proteomics with 0.5 to 5-minute chromatographic gradients at high flow rates (800 µL/min). Using this approach, they quantified 1,937, 2,720, and 4,470 proteins from 5 µg of human cell lysate digests with 0.5-, 1-, and 5-minute LC gradients, respectively. Another group optimized a DIA method that identified >5,000 proteins from 1 μg of HEK293T cell digest in a 5-min gradient using an Orbitrap Exploris 480 MS(*32*). Dia-PASEF has also been successfully employed to fast proteomics analysis with 3-min LC gradient, yielding 5,211 identified proteins from 2 µg cell lysates(*33*). The latest progress in the direction of ultra-fast proteomics method development is the identification of approximately 7,000 proteins with a 5-minute LC gradient on Orbitrap Astral.

In addition to advancements in methodology, the dataset generated in this study introduces several novel aspects and represents a valuable resource for related investigations. Specifically: 1) This study is the first to provide a comprehensive multi-organ proteome-wide analysis of the response mechanisms to ASNase. 2) It integrates large-scale proteomics and metabolomics data for ASNase treatment, offering more detailed and robust evidence at two distinct but interconnected biomolecular levels. 3) While most previous research has focused on the basal levels of ASNS in relation to ASNase efficacy, our dataset provides insights into the quantitative dynamics following drug treatment. 4) Previous studies have primarily examined ASNase treatment for within one or two days. In contrast, our data offer evidence of prolonged drug treatment (up to 8 days), presenting a new perspective on the long-term effects of ASNase.

### Online methods

#### Materials

Water (LC–MS grade, Optima, Cat. No. 10509404), acetonitrile (LC–MS grade, Optima, Cat. No. 10001334), and formic acid (LC–MS grade, Thermo Scientific Pierce, Cat. No. 13454279) were purchased from Fisher Chemicals. HeLa protein digest standard (Thermo Scientific Pierce, Cat. No. 88328) and yeast digest standard (Thermo Scientific Pierce, Cat. No. A47951) were purchased from Thermo Scientific. MassPREP E. coli digest standard (Cat. No. 186003196) was purchased from Waters. Sequence grade trypsin (Cat. No. V5113) were purchased from Promega.

#### Mouse treatment and tissue preparation

The research complies with all relevant ethical regulations: MD Anderson Cancer Center IACUC-approved mouse protocol 00001658-RN02. NSG mice were treated with PBS (vehicle), 20,000 U/kg ASNase^WT^ (Spectrila®) qd for 8 d, or 20,000 U/kg ASNase^Q59L^ qd for 8 d. We previously found the selected doses of ASNase^WT^ and ASNase^Q59L^ to be curative and ineffective, respectively, against a mouse model of ALL(*34*). *Escherichia coli* L-asparaginase II (ASNase) recombinant proteins were produced as described previously(*34, 35*). Treatments were administered intraperitoneally (IP) immediately above the hind leg. Treatment volume was 0.1 mL. Biological samples were collected at t = 0, 1, 3, 5, 8 d (n=3 mice per time point per treatment group). Prior to tissue collection, each mouse was euthanized using a CO2 chamber. The abdominal cavity was opened and perfused with 50 mL pre-cooled phosphate buffered saline. A total of 507 organs were dissected, snap-frozen in liquid nitrogen, and stored at −80°C.

#### Protein extraction and digestion

10 mg of each tissue sample was resuspended in ice-cold protein extraction buffer containing 4 M urea and 50 mM NH_4_HCO_3_. Homogenization was performed using a ceramic bead kit (Bertin Technologies Cat. No.CK28-R) with an automated homogenizer (Precellys 24 with Cryolys adapter, Bertin Technologies) with 2 × 10 s pulses at 5,000 rpm. Homogenates were clarified by centrifugation for 10 min at 10,000 g at 4°C, and protein concentration was determined by the BCA method. Proteins in the solution were denatured by boiling at 95°C for 5 min.

For standard sample preparation workflow, 20 µg protein was diluted to 50 µL with 50 mM NH_4_HCO_3_. Samples were digested by adding 1 µL 400 ng/µL trypsin (Promega) and incubated at 37°C overnight unless otherwise specified. Samples were acidified with formic acid at ∼0.1% (v/v) final concentration by adding 0.5 µL 10% formic acid in water and centrifuged at 10,000 g for 10 min at 4°C. The supernatant was desalted using BioPureSPN Mini PROTO 300 C18 column (The Nest Group, Cat. No. HUM S18V), dried in vacuum and stored at -80°C until analysis.

For high-throughput sample preparation method, 20 µg of protein samples were diluted to 50 µL by adding 50 mM NH4HCO3 in the 96 AFA-TUBE TPX plate (Covaris, Cat. No. 520291) and then add 1 µL 400 ng/µL trypsin (Promega). The sample solutions were incubated with sonication at 37 °C for 4 hours using R230 Focused-Ultrasonicator (Covaris, Cat. No. 500620). Digested samples in 96-well plates were acidified with formic acid at ∼1% (v/v) final concentration by adding 0.5 µL 10% formic acid in water and transferred to Evotip Pure 96-tip racks (Evosep, Cat. No. EV2013) for desalting. The desalted peptide samples were transferred to the Thermo Scientific™ SureSTART™ Level 2 WebSeal™ 96-Well Microtiter Plates (Thermo Scientific, Cat. No. 60180-P210B), dried in vacuum and stored at -80 °C until analysis.

#### LC–MS/MS analysis

LC–MS/MS analysis were performed using Orbitrap Astral mass spectrometer coupled with Vanquish Neo UHPLC system (Thermo Fisher Scientific). The vacuum dried peptide samples were resuspended with 0.1% of formic acid. Peptides were separated using an Evosep endurance column (Evosep, Cat. No. EV1107, 150 μm × 4 cm) at a flow rate of 5 μL/min. Peptides were chromatographically separated using a linear gradient of solvent B (0.1% formic acid in ACN) and solvent A (0.1% formic acid in water). Linear gradients were as follows: from 10% to 38% of solvent B 1.8 min, 38% to 100% of solvent B from 1.8 to 1.9 min, 100% B from 1.9 to 2 min. For the original DIA method on the Orbitrap Astral MS, MS1 spectra were collected in the Orbitrap every 0.6 s at a resolution of 240,000. The full scan range was 380-980 m/z unless otherwise specified. The MS1 normalized AGC target was set to 500% with a maximum injection time of 5 ms. DIA MS2 scans were acquired in the Astral analyzer over a range of 380-980 m/z with a normalized AGC target of 500% and a maximum injection time of 3.5 ms and an HCD collision energy setting of 25%. Window placement optimization was turned on. The isolation window was set at 2 Th without window overlap. For the optimized DIA method, scan range was changed to 495-745 m/z. Isolation window was set at 1 Th without window overlap. The maximum injection time of MS2 was set at 2.5 ms.

#### MS data analysis

Raw files from DIA experiments were analyzed in DIA-NN 1.8.1(*12*). The spectral library was generated from a mouse reference database (UniProt 2024 release, 21,709 sequences) allowing N-term M excision and 1 missed cleavage. The DIA-NN search included the following settings: Protein inference = ‘Genes’, Neural network classifier = ‘Single-pass mode’, Quantification strategy = ‘Robust LC (high precision)’, Cross-run normalization = ‘RT-dependent’, Library Generation = ‘Smart Profiling’ and Speed and RAM usage = ‘Optimal results’. Mass accuracy and MS1 accuracy were set to 0 for automatic inference. ‘No share spectra’, ‘Heuristic protein inference’ and ‘MBR’ were checked. The output results from DIA-NN were filtered for Q.Value <=0.01 and PG.Qvalue <=0.01. DIA-NN R package was used for protein quantitation using MaxLFQ algorithm.

#### Analysis of metabolites

Metabolites profiling was conducted on snap-frozen tissue samples by ultra-high-resolution mass spectrometry. Approximately 20–30 mg of each sample was pulverized in liquid nitrogen and then homogenized with Precellys Tissue Homogenizer.

Metabolites were extracted using ice-cold 0.1% ammonium hydroxide in an 80/20 (v/v) methanol/water solution. The extracts were then centrifuged at 17,000 g for 5 minutes at 4°C, and the supernatants were transferred to clean tubes. The solvents were evaporated under nitrogen to dryness. The dried extracts were reconstituted in deionized water, and 5 μL was injected for analysis by ion chromatography (IC)-mass spectrometry (MS). IC mobile phase A (MPA) was water, and mobile phase B (MPB) was water containing 100 mM KOH. The analysis was performed using a Thermo Scientific Dionex ICS-5000+ capillary IC system with a Thermo IonPac AS11 column (4 µm particle size, 250 × 2 mm). The column compartment was maintained at 30°C, and the autosampler tray was kept at 4°C. The IC flow rate was 350 µL/min with the following gradient conditions: starting at 1 mM KOH, increasing to 35 mM over 25 minutes, further increasing to 99 mM over 39 minutes, and holding at 99 mM for 10 minutes. To enhance desolvation for better sensitivity, methanol was delivered via an external pump and mixed with the post-column eluent using a low dead volume mixing tee. Data acquisition was performed with a Thermo Orbitrap Fusion Tribrid Mass Spectrometer in ESI negative ionization mode with a resolution of 240,000.

For amino acid analysis, samples were extracted using 1 mL of ice-cold 90/10 (v/v) acetonitrile/water containing 0.1% formic acid. The extracts were centrifuged at 17,000g for 5 min at 4°C, and the supernatants were transferred to clean tubes. The solvent was evaporated under nitrogen to dryness. The dried extracts were reconstituted in 90/10 acetonitrile/water with 1% formic acid, and 10 μL was injected for analysis by liquid chromatography (LC)-MS. LC mobile phase A (MPA; weak) was acetonitrile containing 1% formic acid, and mobile phase B (MPB; strong) was water containing 50 mM ammonium formate. The analysis was preformed using Thermo Vanquish LC system equipped with an Imtakt Intrada Amino Acid column (3 µm particle size, 150 × 2.1 mm), maintained at 30°C. The autosampler tray was kept to 4°C. The flow rate was 300 µL/min. The gradient conditions were as follows: initial 15% MPB, increasing to 30% MPB at 20 min, further increasing to 95% MPB at 30 min, holding at 95% MPB for 10 min, and then returning to the initial conditions for equilibration over 10 min. The total run time was 50 min. Data acquisition was performed with a Thermo Orbitrap Fusion Tribrid mass spectrometer in ESI-positive ionization mode at a resolution of 240,000. Raw data files were analyzed using Thermo Trace Finder software. Peaks with a signal-to-noise ratio <3 were considered undetected.

## Supplementary figure legends

**Supplementary Fig. 1.**
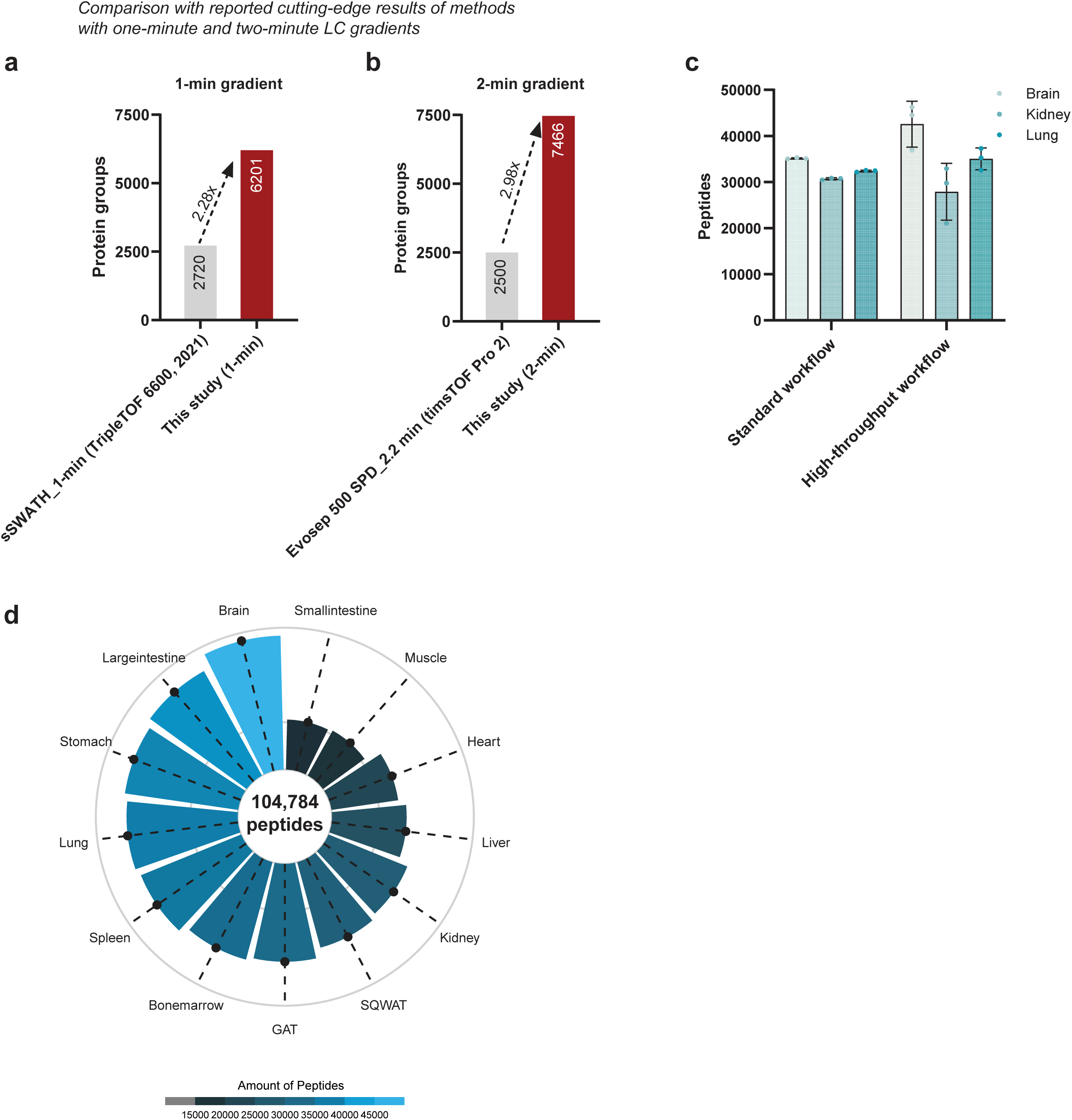
**a,** Comparison of the number of identified protein groups between this study and a reported cutting-edge result of MS methods with one-minute LC gradient. **b**, Comparison of the number of identified protein groups between this study and a reported cutting-edge result of MS methods with two-minute LC gradient. **c,** The number of peptides identified using standard and high-throughput sample preparation workflows from brain, kidney, and lung, respectively. **d,** Overview of the average number of peptides identified in 13 mouse tissues.

**Supplementary Fig. 2.**
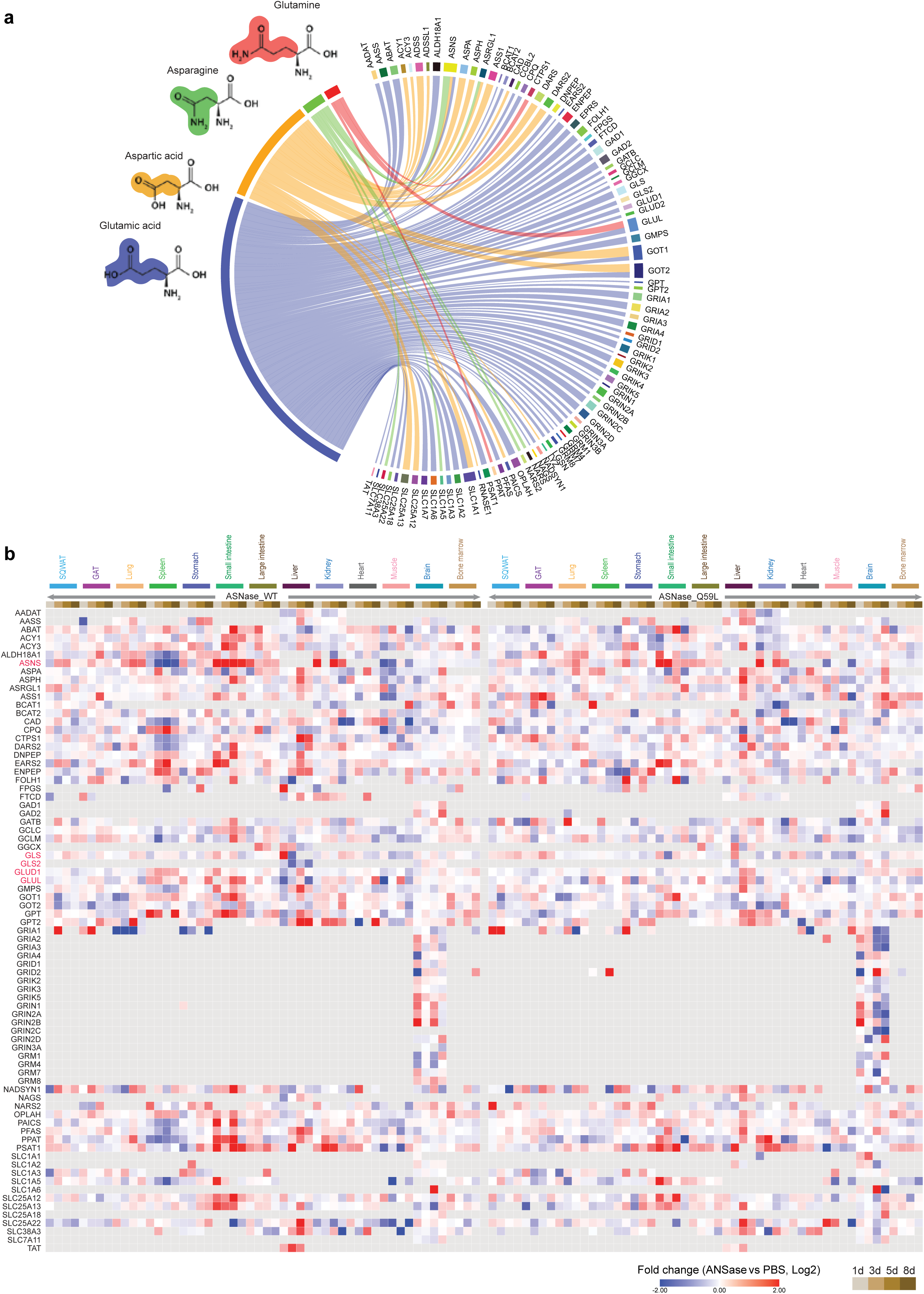
a, Illustration of the binding proteins for four amino acids (ASN, ASP, GLN, and GLU) retrieved from the BioGRID online database (http://thebiogrid.org). **b,** Heatmap overview of the dynamic changes in protein levels of the binding proteins for ASN, ASP, GLN, and GLU following ASNase^WT^ or ASNase^Q59L^ treatment across all the tested tissues. Protein expression levels following ASNase^WT^ or ASNase^Q59L^ treatment were compared to PBS-treated controls, with the Log2 fold change values presented.

**Supplementary Fig. 3.**
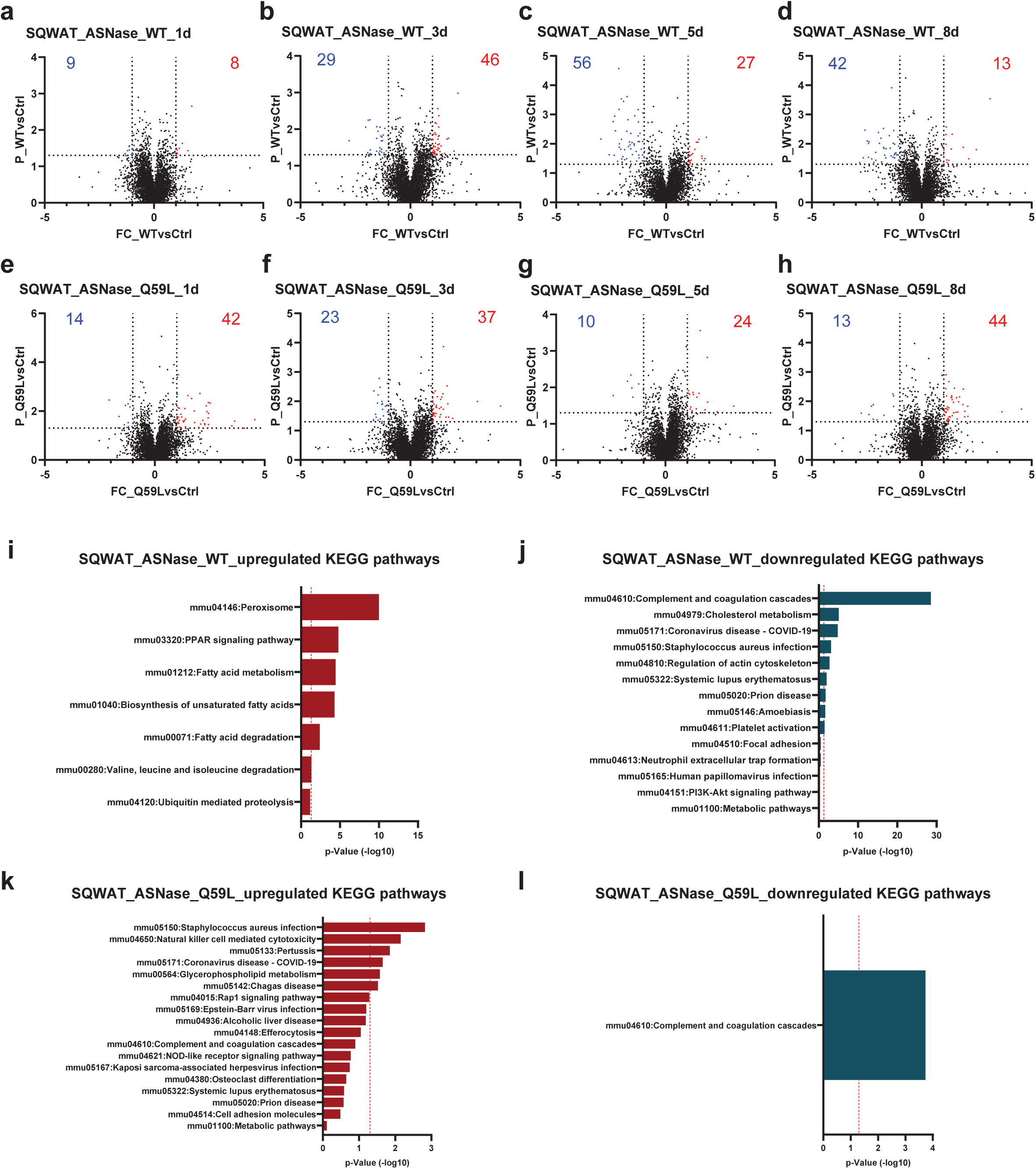
**a-d**, Volcano plots showing differential protein expression levels in SQWAT tissue following ASNase^WT^ treatment at 1d (**a**), 3d (**b**), 5d (**c**), and 8d (**d**). The x-axis represents the Log2 fold changes in protein expression between ASNase^WT^ and PBS-treated controls, while the y-axis indicates the -Log10 of the p-value. Proteins with a fold change greater than 2 and a p-value less than 0.05 are highlighted as significantly upregulated (red) or downregulated (blue). Upregulated or downregulated protein numbers are shown. **e-h**, Similar volcano plots showing differential protein expression levels in SQWAT tissue following ASNase^Q59L^ treatment at 1 day (**e**), 3 days (**f**), 5 days (**g**), and 8 days (**h**). **i**,**j**, KEGG pathway analysis of the upregulated (**i**) or downregulated (**j**) proteins following ASNase^WT^ treatment in the SQWAT tissue. **k**,**l**, KEGG pathway analysis of the upregulated (**k**) or downregulated (**l**) proteins following ASNase^Q59L^ treatment in the SQWAT tissue.

**Supplementary Fig. 4.**
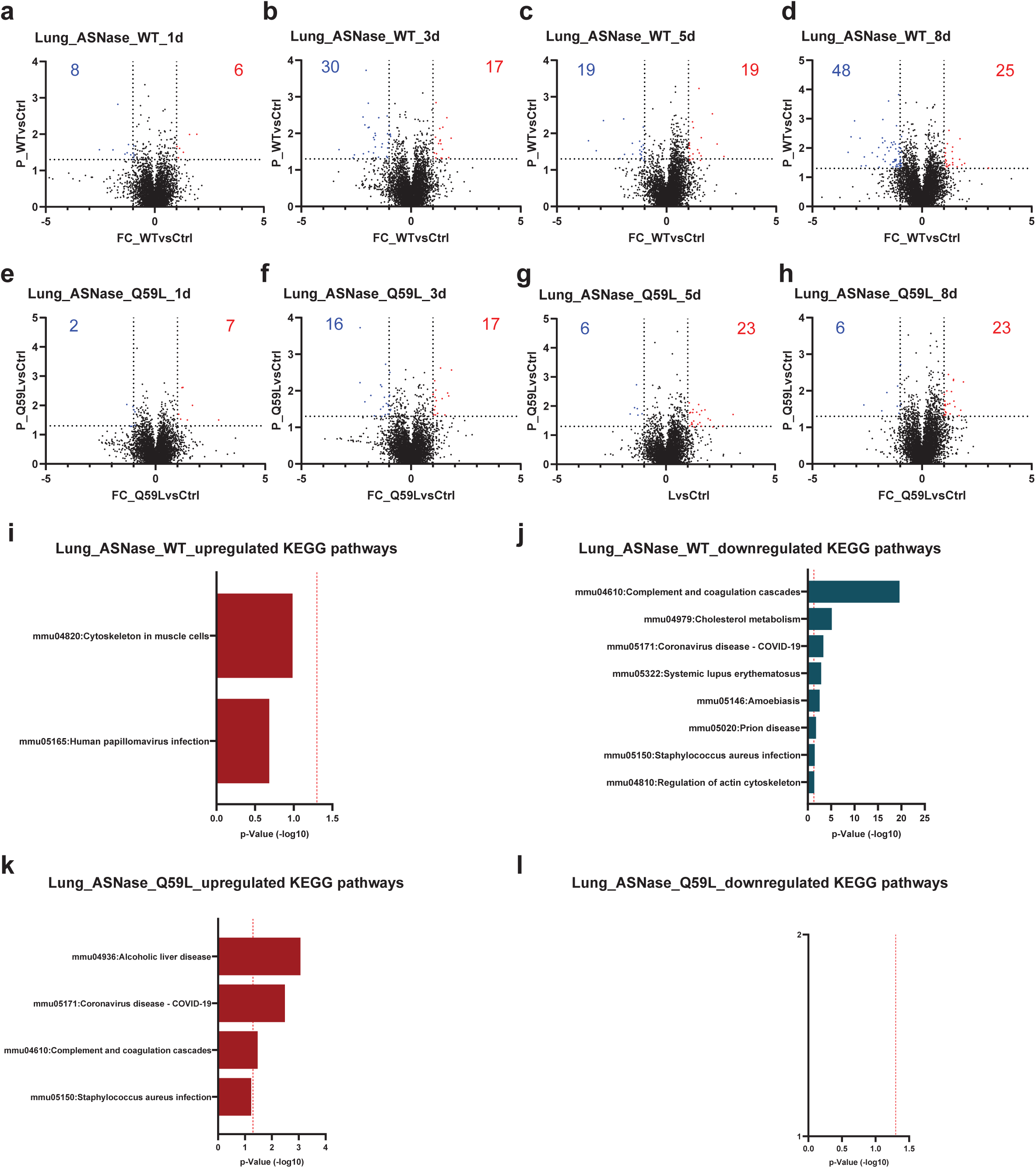
**a-d**, Volcano plots showing differential protein expression levels in lung tissue following ASNase^WT^ treatment at 1d (**a**), 3d (**b**), 5d (**c**), and 8d (**d**). The x-axis represents the Log2 fold changes in protein expression between ASNase^WT^ and PBS-treated controls, while the y-axis indicates the -Log10 of the p-value. Proteins with a fold change greater than 2 and a p-value less than 0.05 are highlighted as significantly upregulated (red) or downregulated (blue). Upregulated or downregulated protein numbers are shown. **e-h**, Similar volcano plots showing differential protein expression levels in lung tissue following ASNase^Q59L^ treatment at 1 day (**e**), 3 days (**f**), 5 days (**g**), and 8 days (**h**). **i**,**j**, KEGG pathway analysis of the upregulated (**i**) or downregulated (**j**) proteins following ASNase^WT^ treatment in the lung tissue. **k**,**l**, KEGG pathway analysis of the upregulated (**k**) or downregulated (**l**) proteins following ASNase^Q59L^ treatment in the lung tissue.

**Supplementary Fig. 5.**
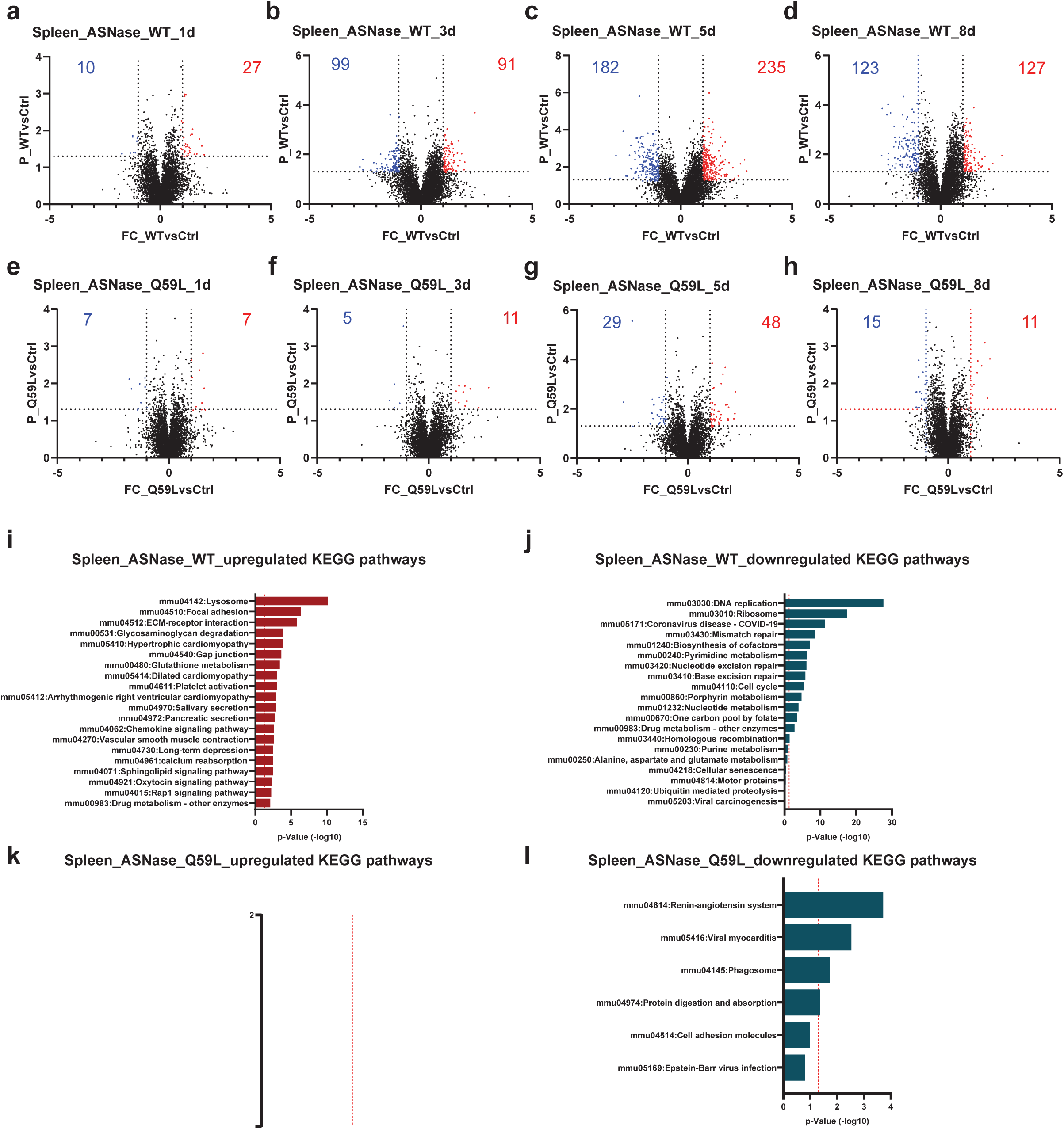
**a-d**, Volcano plots showing differential protein expression levels in spleen tissue following ASNase^WT^ treatment at 1d (**a**), 3d (**b**), 5d (**c**), and 8d (**d**). The x-axis represents the Log2 fold changes in protein expression between ASNase^WT^ and PBS-treated controls, while the y-axis indicates the -Log10 of the p-value. Proteins with a fold change greater than 2 and a p-value less than 0.05 are highlighted as significantly upregulated (red) or downregulated (blue). Upregulated or downregulated protein numbers are shown. **e-h**, Similar volcano plots showing differential protein expression levels in spleen tissue following ASNase^Q59L^ treatment at 1 day (**e**), 3 days (**f**), 5 days (**g**), and 8 days (**h**). **i**,**j**, KEGG pathway analysis of the upregulated (**i**) or downregulated (**j**) proteins following ASNase^WT^ treatment in the spleen tissue. **k**,**l**, KEGG pathway analysis of the upregulated (**k**) or downregulated (**l**) proteins following ASNase^Q59L^ treatment in the spleen tissue.

**Supplementary Fig. 6.**
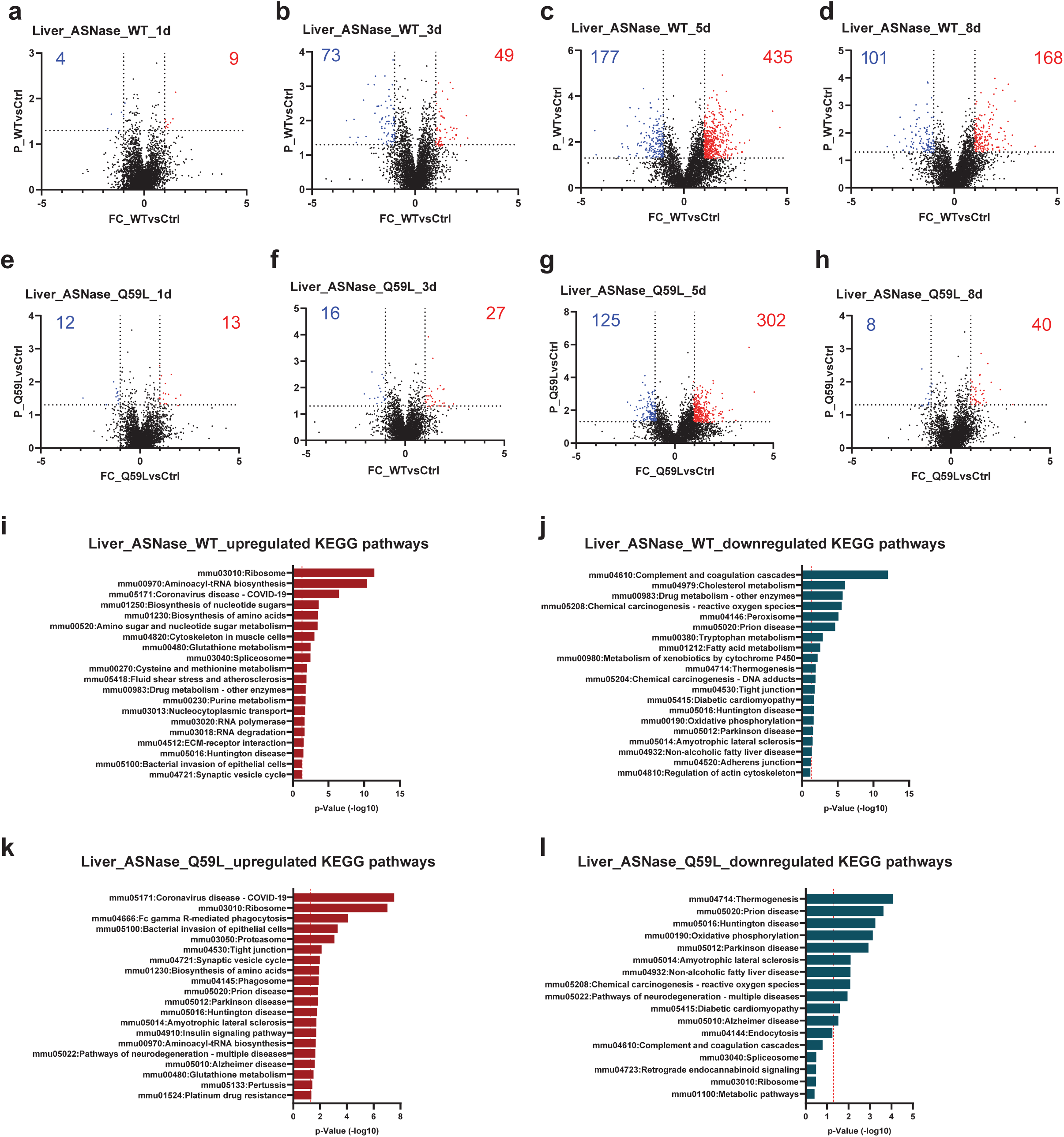
**a-d**, Volcano plots showing differential protein expression levels in liver tissue following ASNase^WT^ treatment at 1d (**a**), 3d (**b**), 5d (**c**), and 8d (**d**). The x-axis represents the Log2 fold changes in protein expression between ASNase^WT^ and PBS-treated controls, while the y-axis indicates the -Log10 of the p-value. Proteins with a fold change greater than 2 and a p-value less than 0.05 are highlighted as significantly upregulated (red) or downregulated (blue). Upregulated or downregulated protein numbers are shown. **e-h**, Similar volcano plots showing differential protein expression levels in liver tissue following ASNase^Q59L^ treatment at 1 day (**e**), 3 days (**f**), 5 days (**g**), and 8 days (**h**). **i**,**j**, KEGG pathway analysis of the upregulated (**i**) or downregulated (**j**) proteins following ASNase^WT^ treatment in the liver tissue. **k**,**l**, KEGG pathway analysis of the upregulated (**k**) or downregulated (**l**) proteins following ASNase^Q59L^ treatment in the liver tissue.

**Supplementary Fig. 7.**
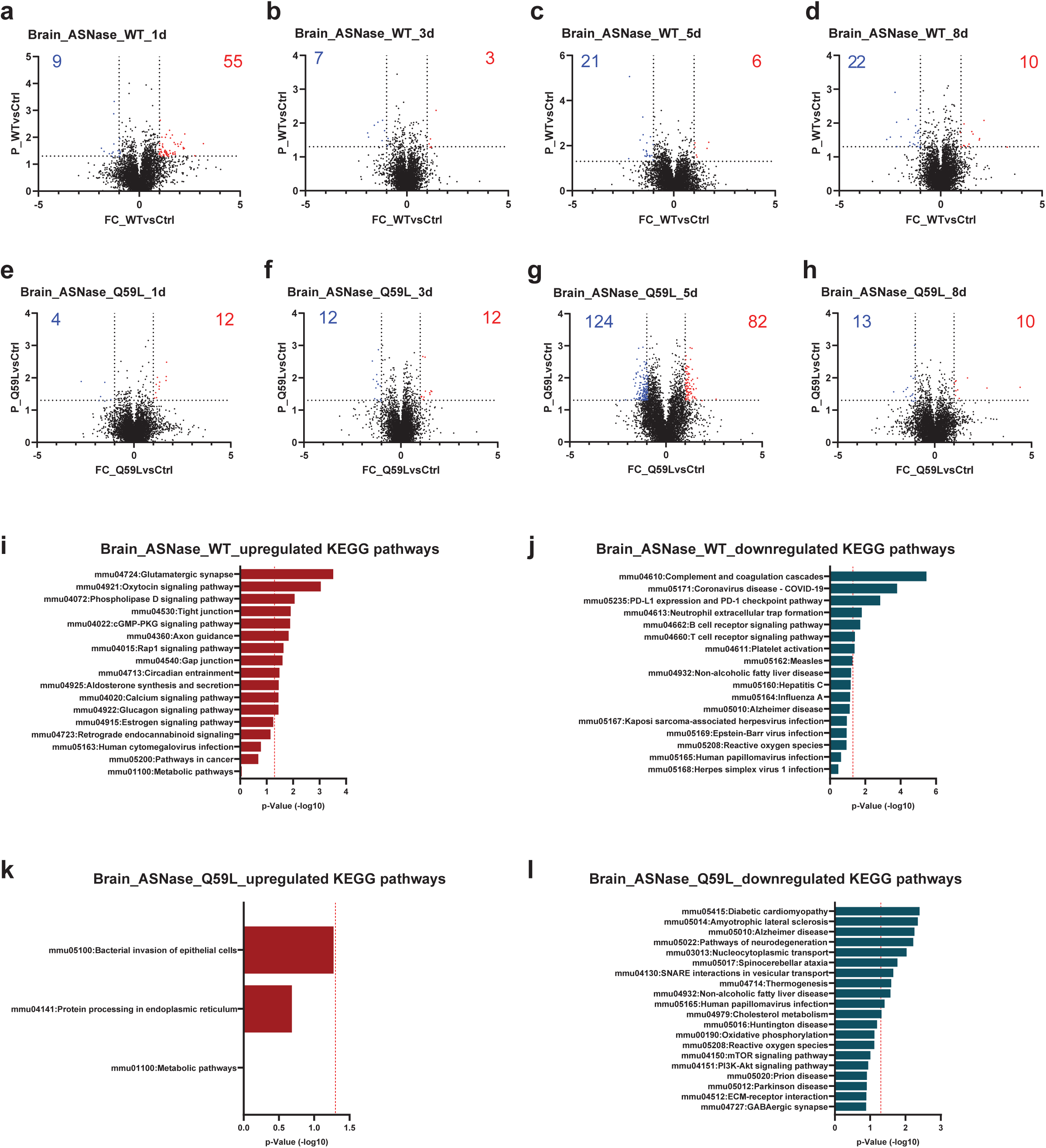
**a-d**, Volcano plots showing differential protein expression levels in brain tissue following ASNase^WT^ treatment at 1d (**a**), 3d (**b**), 5d (**c**), and 8d (**d**). The x-axis represents the Log2 fold changes in protein expression between ASNase^WT^ and PBS-treated controls, while the y-axis indicates the -Log10 of the p-value. Proteins with a fold change greater than 2 and a p-value less than 0.05 are highlighted as significantly upregulated (red) or downregulated (blue). Upregulated or downregulated protein numbers are shown. **e-h**, Similar volcano plots showing differential protein expression levels in brain tissue following ASNase^Q59L^ treatment at 1 day (**e**), 3 days (**f**), 5 days (**g**), and 8 days (**h**). **i**,**j**, KEGG pathway analysis of the upregulated (**i**) or downregulated (**j**) proteins following ASNase^WT^ treatment in the brain tissue. **k**,**l**, KEGG pathway analysis of the upregulated (**k**) or downregulated (**l**) proteins following ASNase^Q59L^ treatment in the brain tissue.

**Supplementary Fig. 8.**
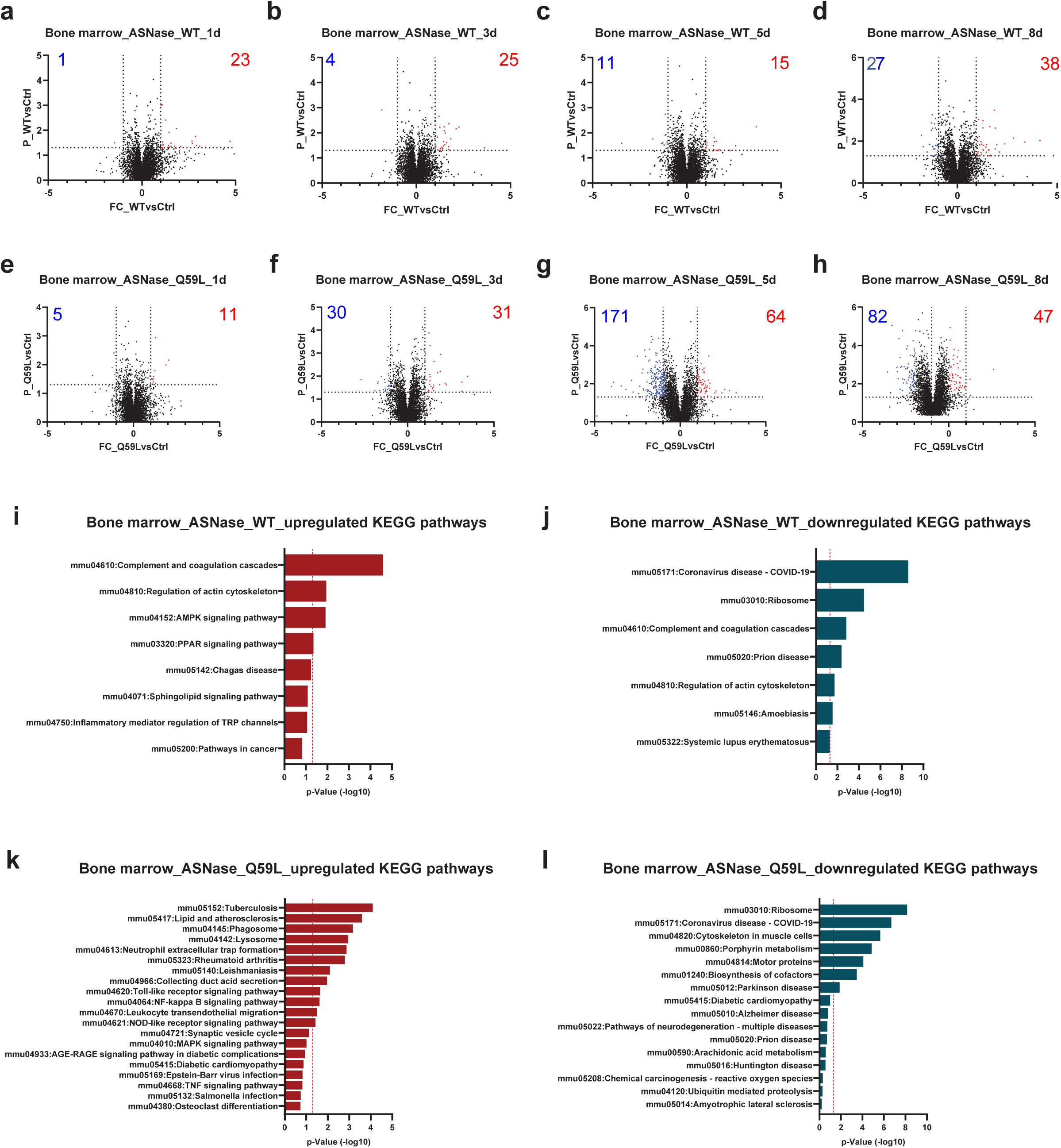
**a-d**, Volcano plots showing differential protein expression levels in bone marrow tissue following ASNase^WT^ treatment at 1d (**a**), 3d (**b**), 5d (**c**), and 8d (**d**). The x-axis represents the Log2 fold changes in protein expression between ASNase^WT^ and PBS-treated controls, while the y-axis indicates the -Log10 of the p-value. Proteins with a fold change greater than 2 and a p-value less than 0.05 are highlighted as significantly upregulated (red) or downregulated (blue). Upregulated or downregulated protein numbers are shown. **e-h**, Similar volcano plots showing differential protein expression levels in bone marrow tissue following ASNase^Q59L^ treatment at 1 day (**e**), 3 days (**f**), 5 days (**g**), and 8 days (**h**). **i**,**j**, KEGG pathway analysis of the upregulated (**i**) or downregulated (**j**) proteins following ASNase^WT^ treatment in the bone marrow tissue. **k**,**l**, KEGG pathway analysis of the upregulated (**k**) or downregulated (**l**) proteins following ASNase^Q59L^ treatment in the bone marrow tissue.

**Supplementary Fig. 9.**
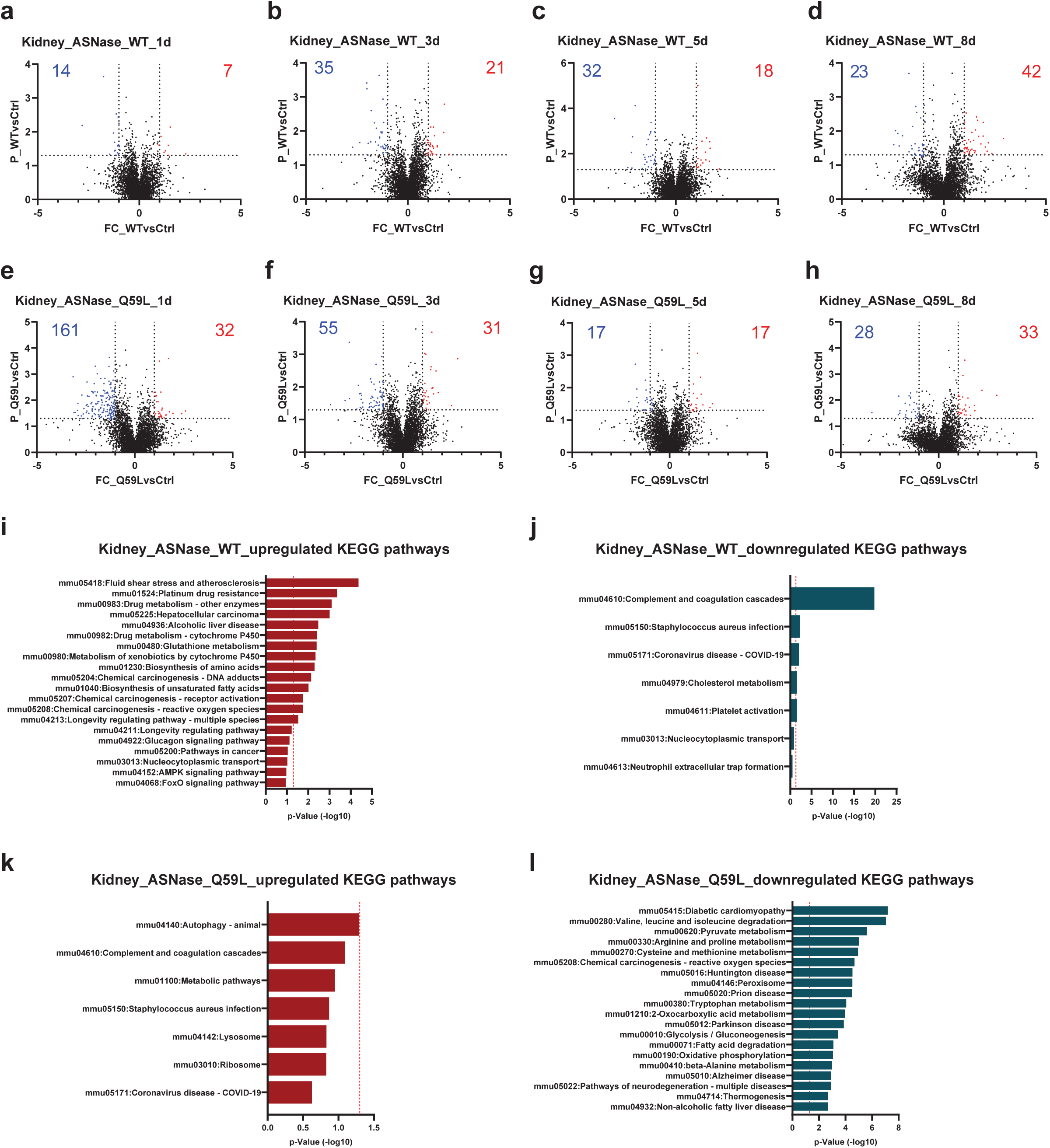
**a-d**, Volcano plots showing differential protein expression levels in kidney tissue following ASNase^WT^ treatment at 1d (**a**), 3d (**b**), 5d (**c**), and 8d (**d**). The x-axis represents the Log2 fold changes in protein expression between ASNase^WT^ and PBS-treated controls, while the y-axis indicates the -Log10 of the p-value. Proteins with a fold change greater than 2 and a p-value less than 0.05 are highlighted as significantly upregulated (red) or downregulated (blue). Upregulated or downregulated protein numbers are shown. **e-h**, Similar volcano plots showing differential protein expression levels in kidney tissue following ASNase^Q59L^ treatment at 1 day (**e**), 3 days (**f**), 5 days (**g**), and 8 days (**h**). **i**,**j**, KEGG pathway analysis of the upregulated (**i**) or downregulated (**j**) proteins following ASNase^WT^ treatment in the kidney tissue. **k**,**l**, KEGG pathway analysis of the upregulated (**k**) or downregulated (**l**) proteins following ASNase^Q59L^ treatment in the kidney tissue.

**Supplementary Fig. 10.**
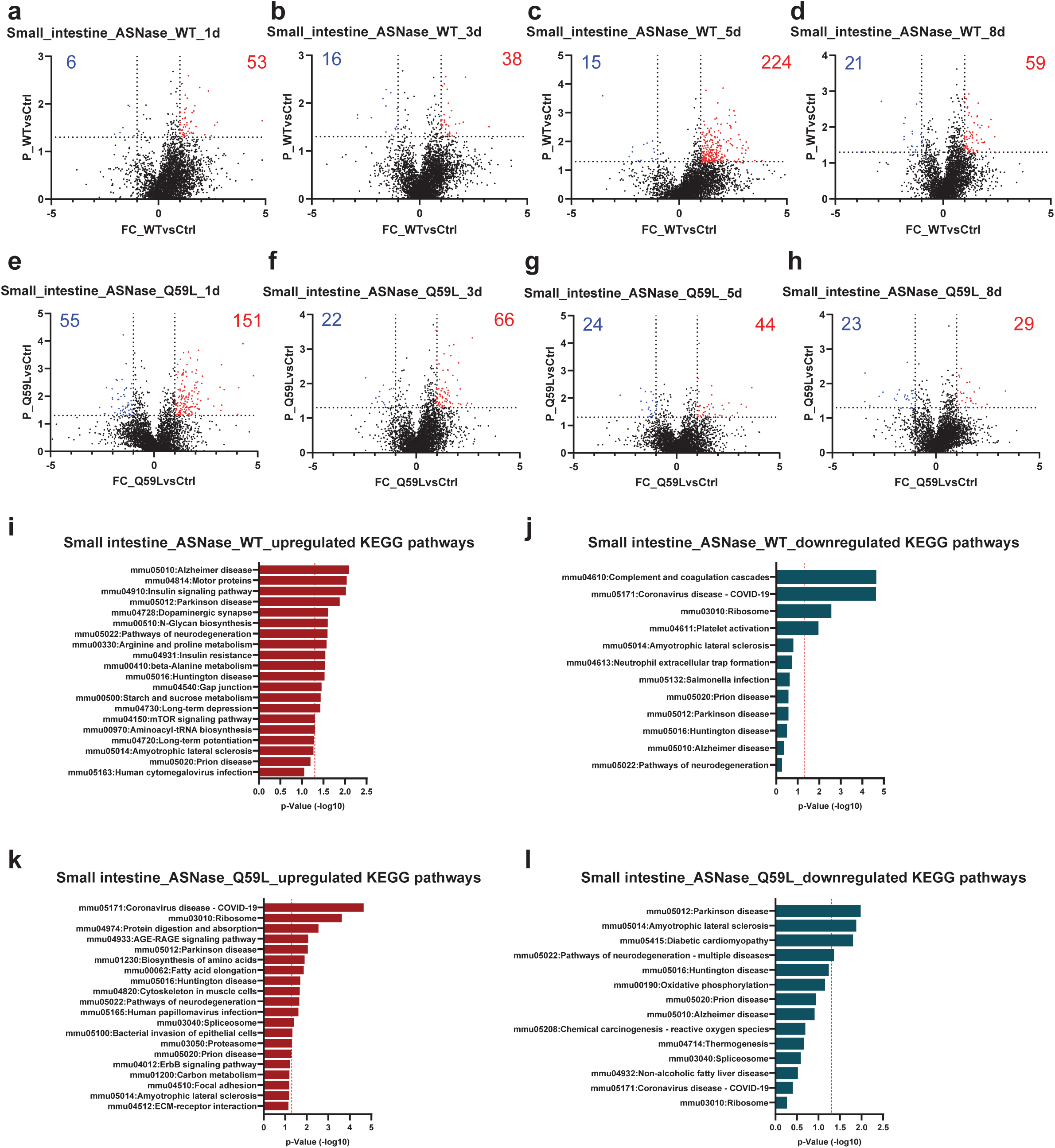
**a-d**, Volcano plots showing differential protein expression levels in small intestine tissue following ASNase^WT^ treatment at 1d (**a**), 3d (**b**), 5d (**c**), and 8d (**d**). The x-axis represents the Log2 fold changes in protein expression between ASNase^WT^ and PBS-treated controls, while the y-axis indicates the -Log10 of the p-value. Proteins with a fold change greater than 2 and a p-value less than 0.05 are highlighted as significantly upregulated (red) or downregulated (blue). Upregulated or downregulated protein numbers are shown. **e-h**, Similar volcano plots showing differential protein expression levels in small intestine tissue following ASNase^Q59L^ treatment at 1 day (**e**), 3 days (**f**), 5 days (**g**), and 8 days (**h**). **i**,**j**, KEGG pathway analysis of the upregulated (**i**) or downregulated (**j**) proteins following ASNase^WT^ treatment in the small intestine tissue. **k**,**l**, KEGG pathway analysis of the upregulated (**k**) or downregulated (**l**) proteins following ASNase^Q59L^ treatment in the small intestine tissue.

**Supplementary Fig. 11.**
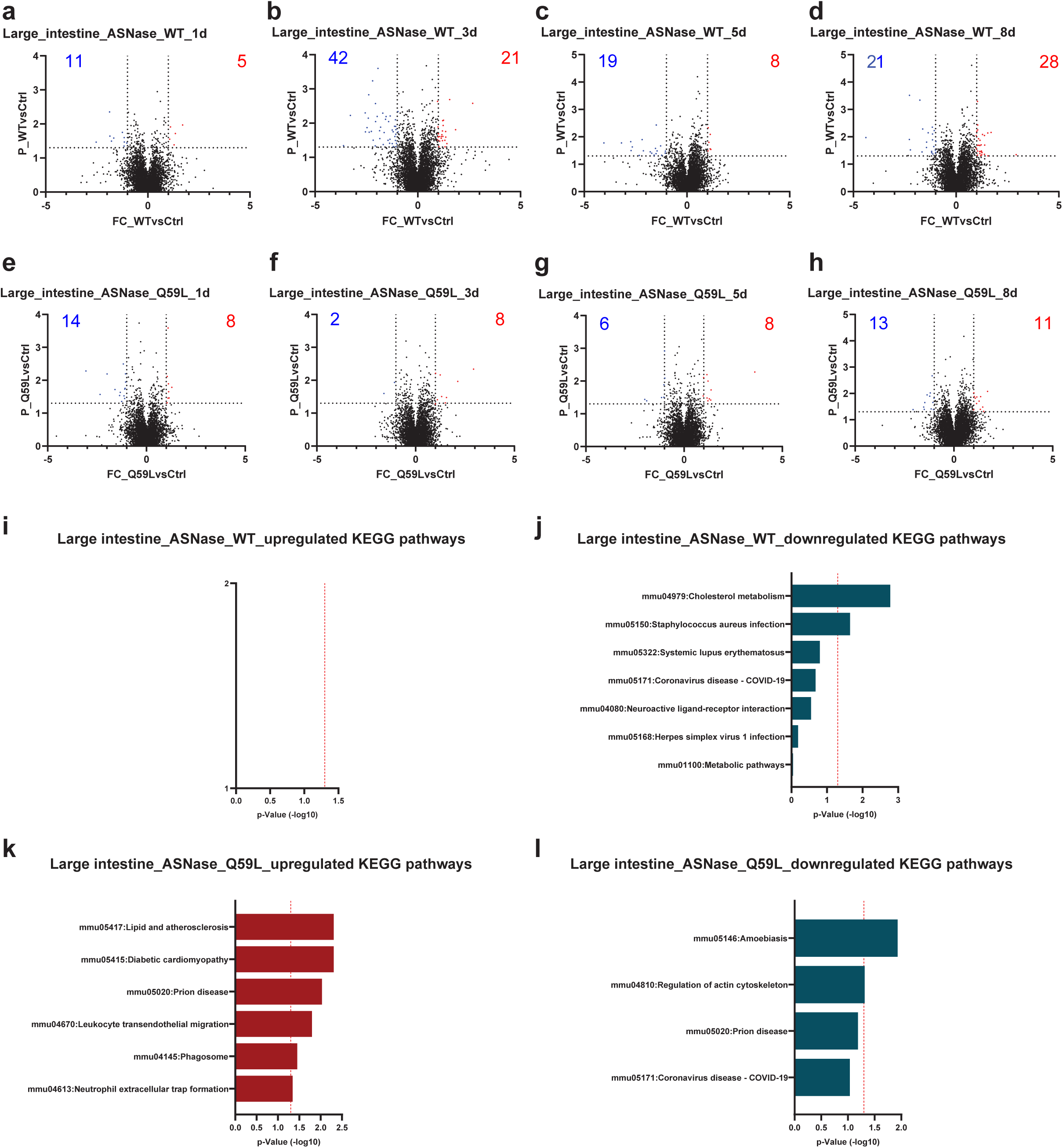
**a-d**, Volcano plots showing differential protein expression levels in large intestine tissue following ASNase^WT^ treatment at 1d (**a**), 3d (**b**), 5d (**c**), and 8d (**d**). The x-axis represents the Log2 fold changes in protein expression between ASNase^WT^ and PBS-treated controls, while the y-axis indicates the -Log10 of the p-value. Proteins with a fold change greater than 2 and a p-value less than 0.05 are highlighted as significantly upregulated (red) or downregulated (blue). Upregulated or downregulated protein numbers are shown. **e-h**, Similar volcano plots showing differential protein expression levels in large intestine tissue following ASNase^Q59L^ treatment at 1 day (**e**), 3 days (**f**), 5 days (**g**), and 8 days (**h**). **i**,**j**, KEGG pathway analysis of the upregulated (**i**) or downregulated (**j**) proteins following ASNase^WT^ treatment in the large intestine tissue. **k**,**l**, KEGG pathway analysis of the upregulated (**k**) or downregulated (**l**) proteins following ASNase^Q59L^ treatment in the large intestine tissue.

**Supplementary Fig.12.**
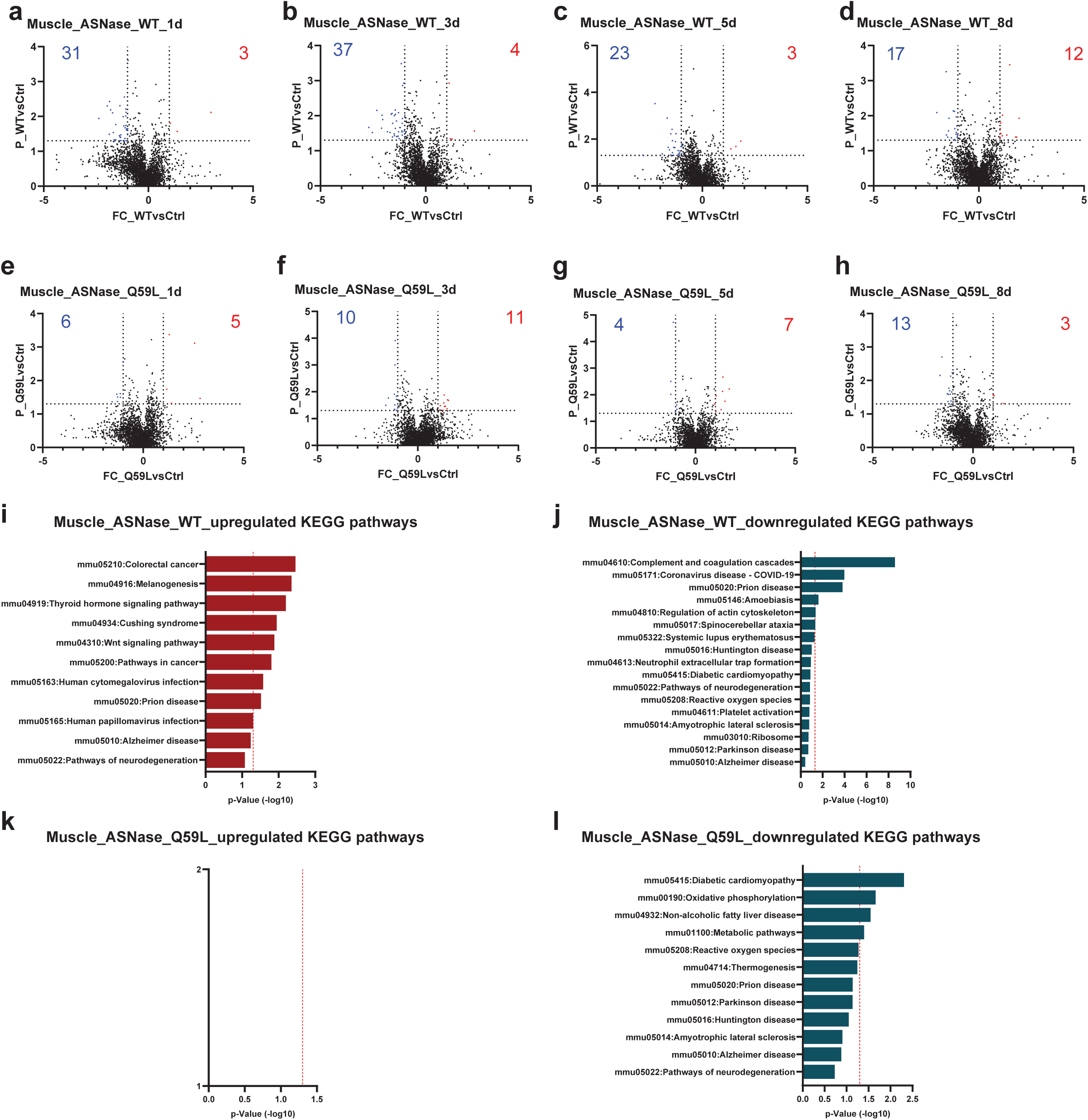
**a-d**, Volcano plots showing differential protein expression levels in muscle tissue following ASNase^WT^ treatment at 1d (**a**), 3d (**b**), 5d (**c**), and 8d (**d**). The x-axis represents the Log2 fold changes in protein expression between ASNase^WT^ and PBS-treated controls, while the y-axis indicates the -Log10 of the p-value. Proteins with a fold change greater than 2 and a p-value less than 0.05 are highlighted as significantly upregulated (red) or downregulated (blue). Upregulated or downregulated protein numbers are shown. **e-h**, Similar volcano plots showing differential protein expression levels in muscle tissue following ASNase^Q59L^ treatment at 1 day (**e**), 3 days (**f**), 5 days (**g**), and 8 days (**h**). **i**,**j**, KEGG pathway analysis of the upregulated (**i**) or downregulated (**j**) proteins following ASNase^WT^ treatment in the muscle tissue. **k**,**l**, KEGG pathway analysis of the upregulated (**k**) or downregulated (**l**) proteins following ASNase^Q59L^ treatment in the muscle tissue.

**Supplementary Fig.13.**
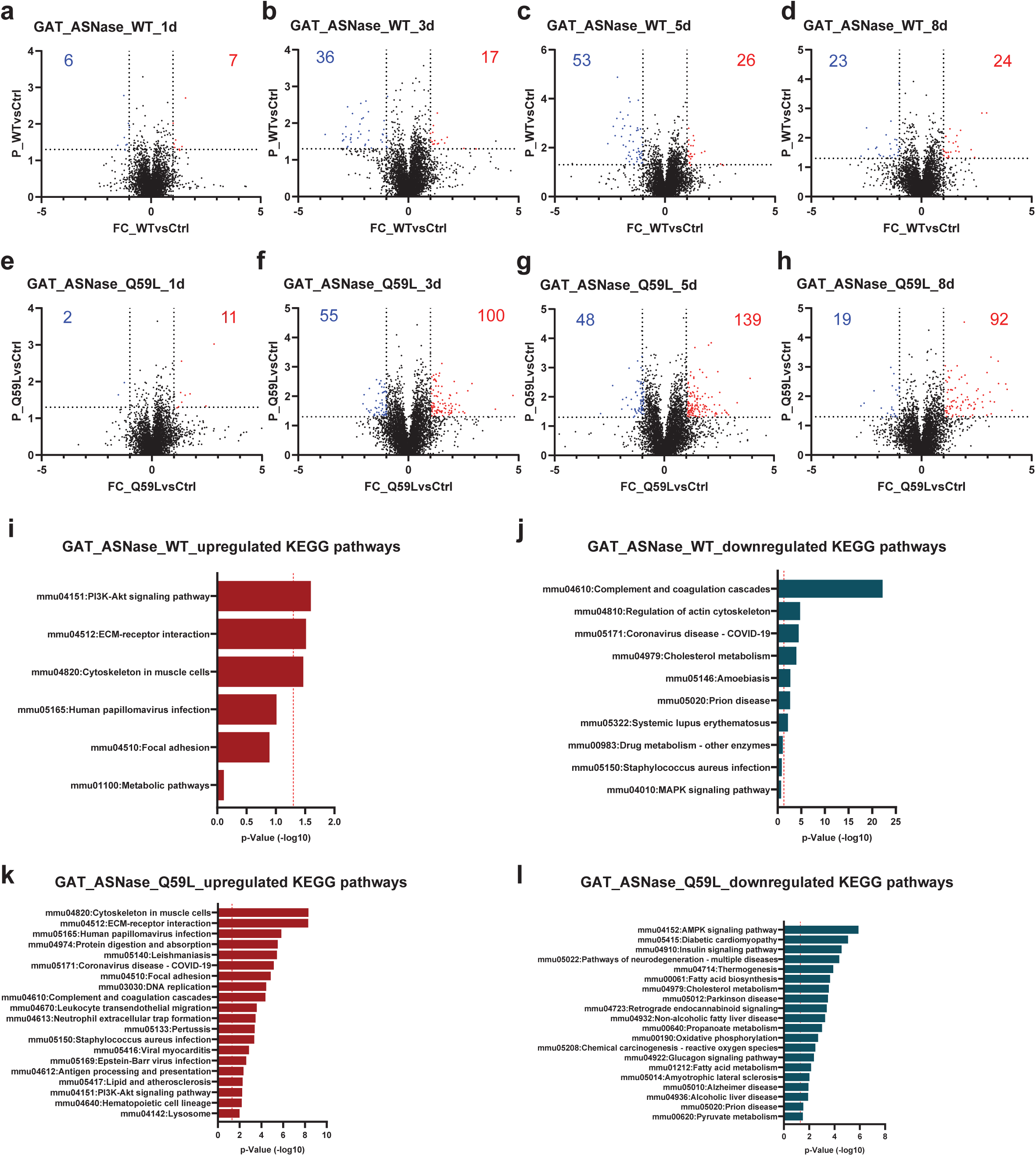
**a-d**, Volcano plots showing differential protein expression levels in GAT tissue following ASNase^WT^ treatment at 1d (**a**), 3d (**b**), 5d (**c**), and 8d (**d**). The x-axis represents the Log2 fold changes in protein expression between ASNase^WT^ and PBS-treated controls, while the y-axis indicates the -Log10 of the p-value. Proteins with a fold change greater than 2 and a p-value less than 0.05 are highlighted as significantly upregulated (red) or downregulated (blue). Upregulated or downregulated protein numbers are shown. **e-h**, Similar volcano plots showing differential protein expression levels in GAT tissue following ASNase^Q59L^ treatment at 1 day (**e**), 3 days (**f**), 5 days (**g**), and 8 days (**h**). **i**,**j**, KEGG pathway analysis of the upregulated (**i**) or downregulated (**j**) proteins following ASNase^WT^ treatment in the GAT tissue. **k**,**l**, KEGG pathway analysis of the upregulated (**k**) or downregulated (**l**) proteins following ASNase^Q59L^ treatment in the GAT tissue.

**Supplementary Fig.14.**
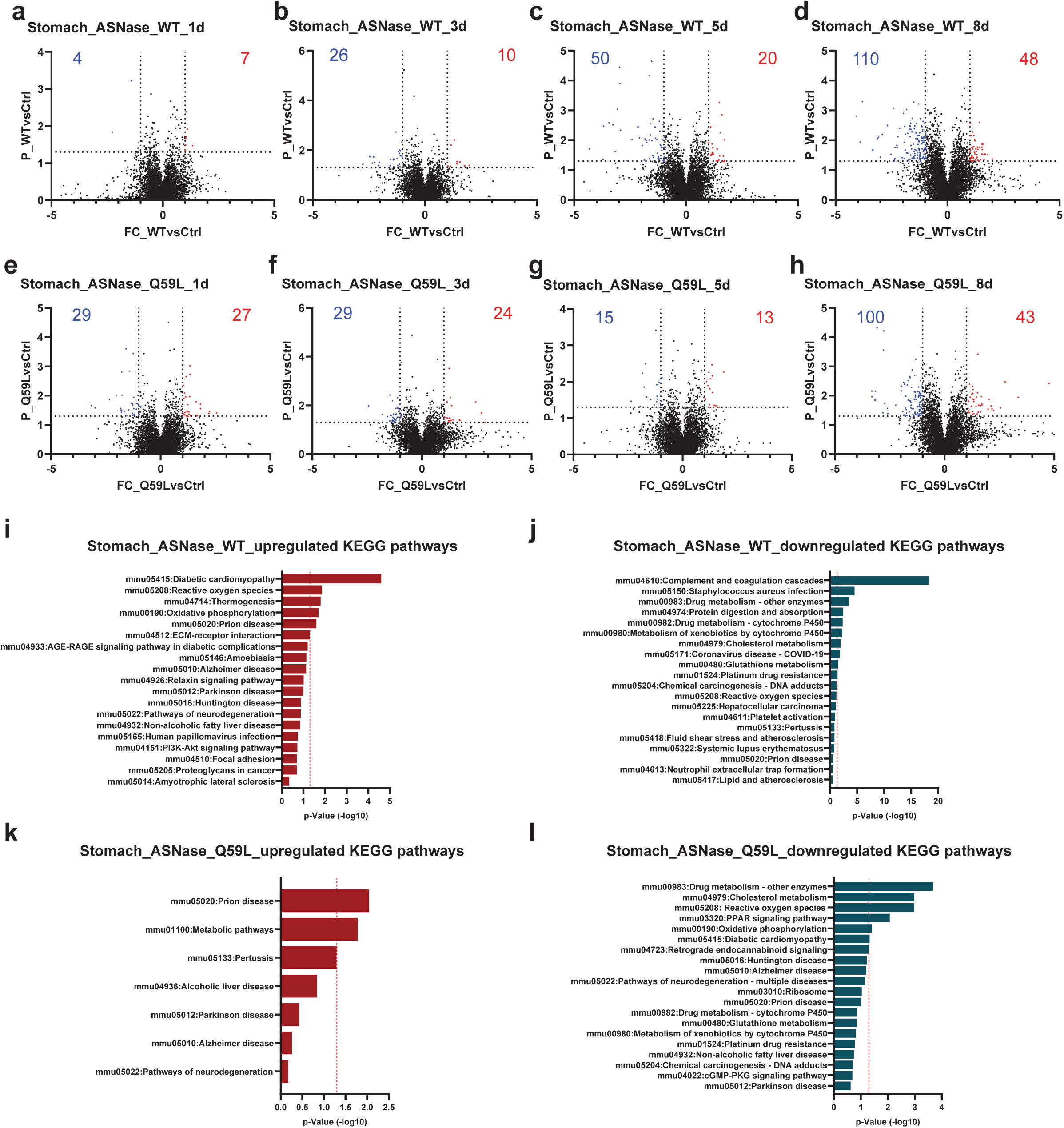
**a-d**, Volcano plots showing differential protein expression levels in stomach tissue following ASNase^WT^ treatment at 1d (**a**), 3d (**b**), 5d (**c**), and 8d (**d**). The x-axis represents the Log2 fold changes in protein expression between ASNase^WT^ and PBS-treated controls, while the y-axis indicates the -Log10 of the p-value. Proteins with a fold change greater than 2 and a p-value less than 0.05 are highlighted as significantly upregulated (red) or downregulated (blue). Upregulated or downregulated protein numbers are shown. **e-h**, Similar volcano plots showing differential protein expression levels in stomach tissue following ASNase^Q59L^ treatment at 1 day (**e**), 3 days (**f**), 5 days (**g**), and 8 days (**h**). **i**,**j**, KEGG pathway analysis of the upregulated (**i**) or downregulated (**j**) proteins following ASNase^WT^ treatment in the stomach tissue. **k**,**l**, KEGG pathway analysis of the upregulated (**k**) or downregulated (**l**) proteins following ASNase^Q59L^ treatment in the stomach tissue.

**Supplementary Fig.15.**
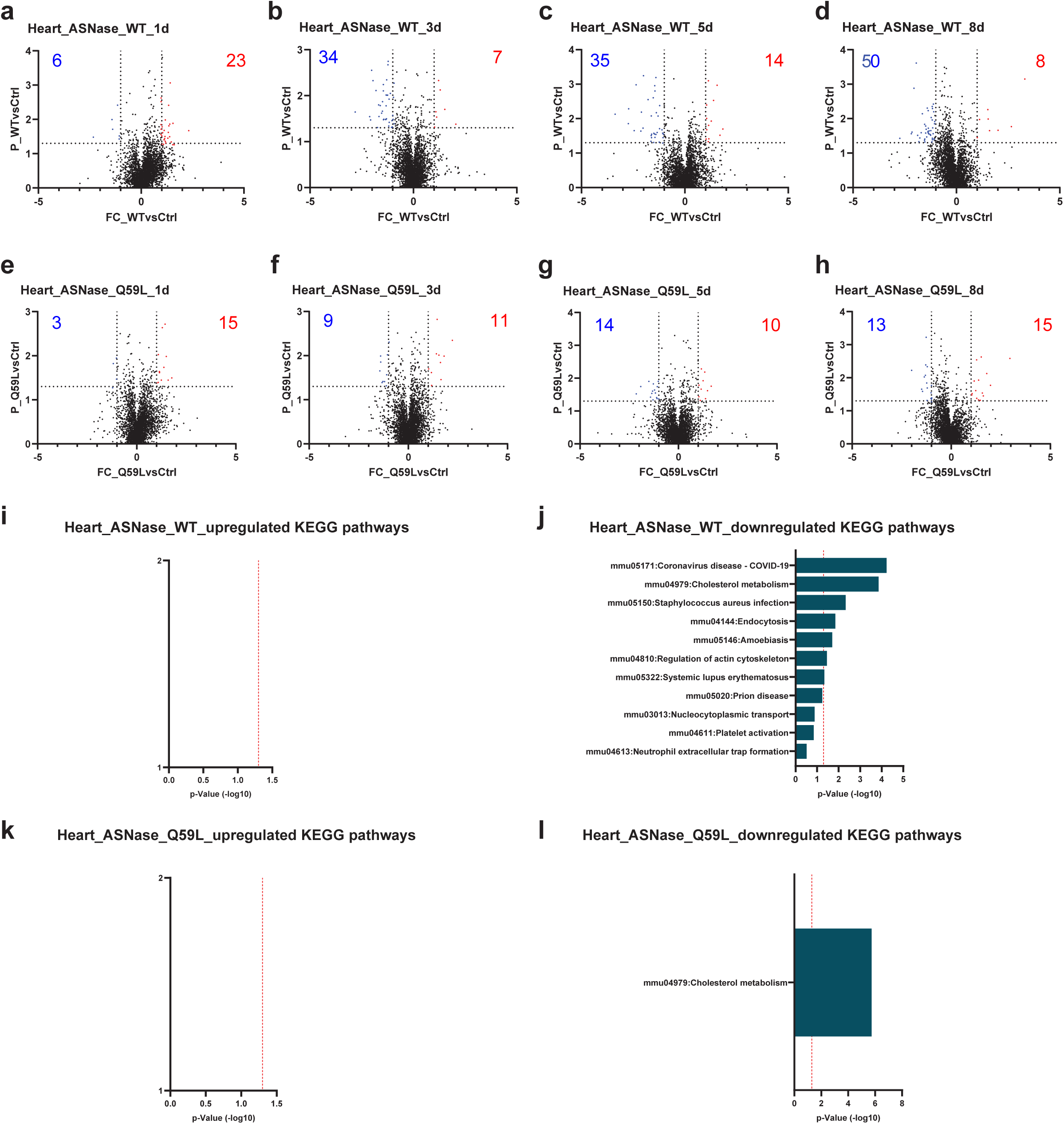
**a-d**, Volcano plots showing differential protein expression levels in heart tissue following ASNase^WT^ treatment at 1d (**a**), 3d (**b**), 5d (**c**), and 8d (**d**). The x-axis represents the Log2 fold changes in protein expression between ASNase^WT^ and PBS-treated controls, while the y-axis indicates the -Log10 of the p-value. Proteins with a fold change greater than 2 and a p-value less than 0.05 are highlighted as significantly upregulated (red) or downregulated (blue). Upregulated or downregulated protein numbers are shown. **e-h**, Similar volcano plots showing differential protein expression levels in heart tissue following ASNase^Q59L^ treatment at 1 day (**e**), 3 days (**f**), 5 days (**g**), and 8 days (**h**). **i**,**j**, KEGG pathway analysis of the upregulated (**i**) or downregulated (**j**) proteins following ASNase^WT^ treatment in the heart tissue. **k**,**l**, KEGG pathway analysis of the upregulated (**k**) or downregulated (**l**) proteins following ASNase^Q59L^ treatment in the heart tissue.

**Supplementary Fig. 16.**
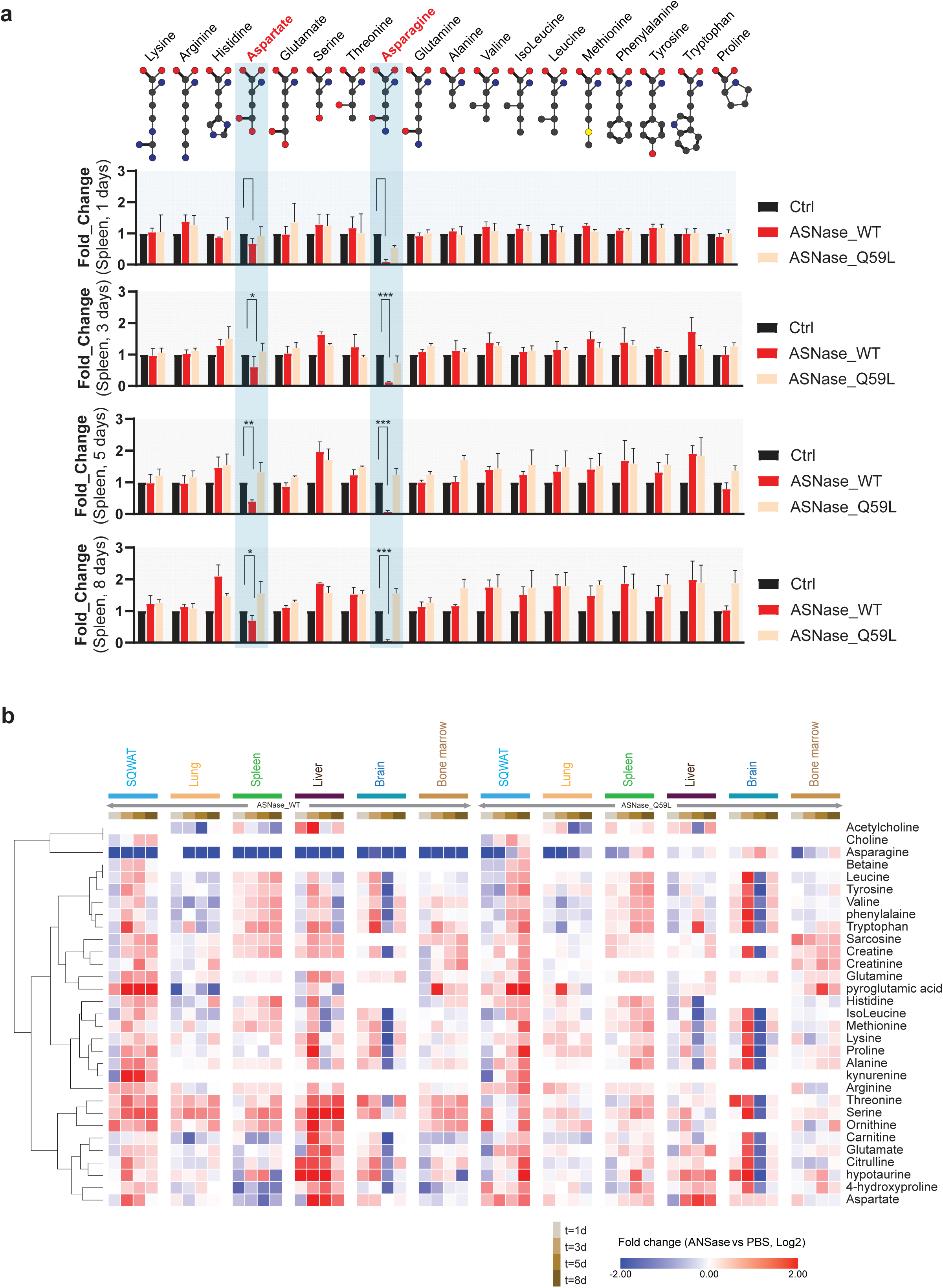
a, The dynamic regulation of indicated amino acid levels following 1, 3, 5, 8 days treatment with ASNase^WT^ or ASNase^Q59L^ treatment in the spleen. **b,** The dynamic regulation of indicated amino acid levels following 1, 3, 5, 8 days treatment with ASNase^WT^ or ASNase^Q59L^ treatment across all tested tissues.

**Supplementary Fig. 17.**
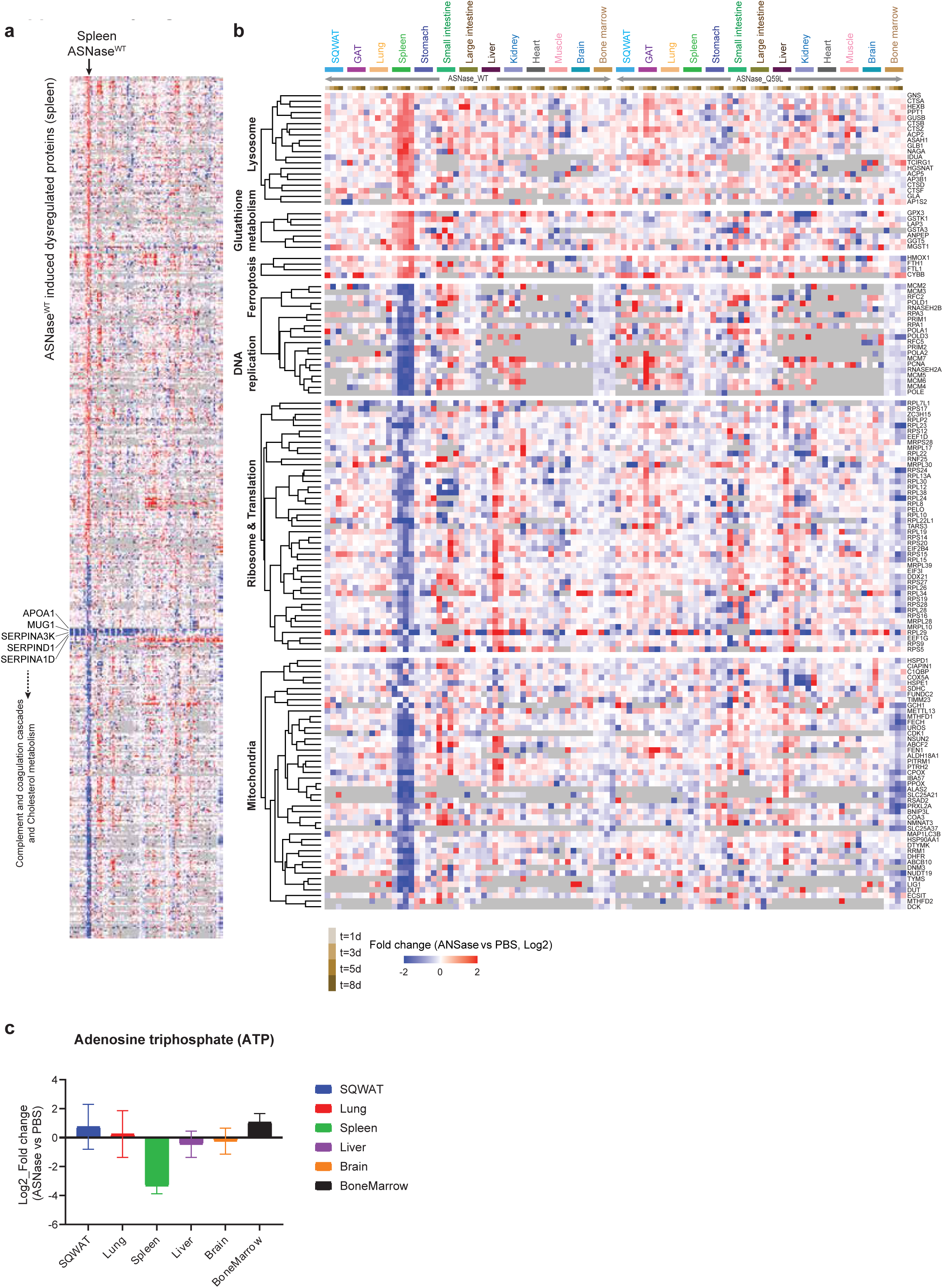
**a**, Proteins that were upregulated or downregulated in the spleen following ASNase^WT^ treatment were categorized. A heatmap was used to provide a global overview of the dynamic changes in these proteins across all tested tissues. **b**, An enlarged view of (**a**) highlighted proteins involved in lysosome, glutathione metabolism, ferroptosis, DNA replication, ribosome and translation, and mitochondria.

**Supplementary Fig. 18.**
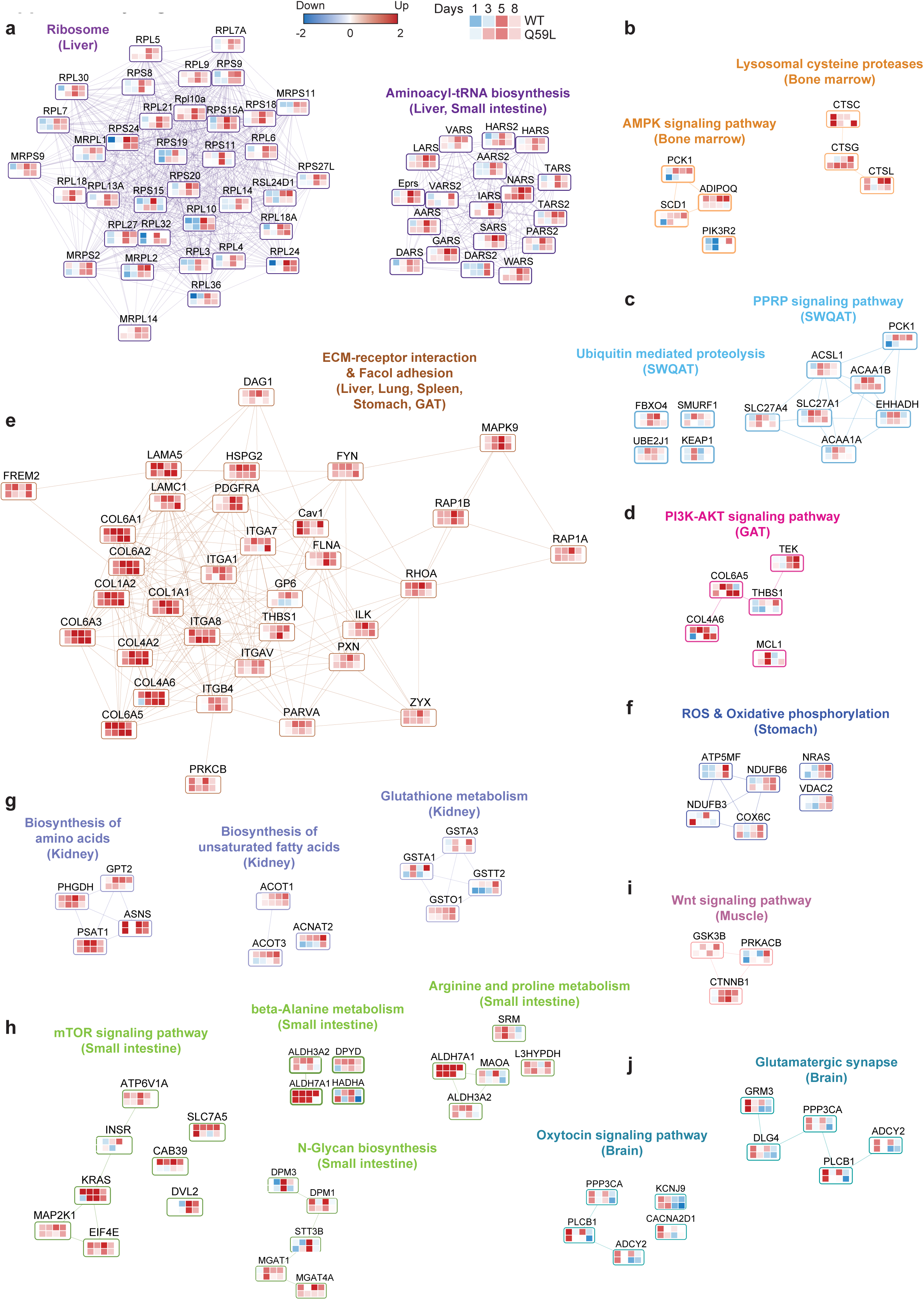
Landscape of tissue specific dysregulation pathways following ASNase^WT^ treatment. **a,** Proteins involved in the up-regulated ribosomal proteins and aminoacyl-tRNA biosynthesis in the liver. **b,** Proteins involved in the up-regulated proteins involved in AMPK signaling pathway and lysosomal cysteine proteases in the bone marrow. **c,** Proteins involved in the up-regulated proteins involved in the PPRP signaling pathway and the ubiquitin mediated proteolysis in the SWQAT. **d,** Proteins involved in the up-regulated proteins involved in PI3K-AKT signaling pathway in the GAT. **e,** Proteins involved in the up-regulated proteins involved in the ECM-receptor interaction and focal adhesion in multiple tissues including liver, lung, spleen, stomach, and GAT. For visualization, the maximal log2 ratios (ASNase^WT^ vs PBS) from liver, lung, spleen, stomach, or GAT were adopted for the heatmaps. **f,** Proteins involved in the up-regulated proteins involved in ROS and oxidative phosphorylation pathway in the stomach. **g,** Proteins involved in the up-regulated proteins involved in biosynthesis of amino acids pathway, biosynthesis of unsaturated fatty acids pathway, and glutathione metabolism pathway in the kidney. **h,** Proteins involved in the up-regulated proteins involved in mTOR signaling pathway, beta-Alanine metabolism pathway, arginine and proline metabolism pathway and N-glycan biosynthesis pathway in the small intestine. **j,** Proteins involved in the up-regulated proteins involved in glutamatergic synapse and oxytocin signaling pathway in the brain. The maximal log2 ratios (ASNase^WT^ vs PBS) from indicated tissues were adopted for the heatmaps unless otherwise specified.

**Supplementary Fig. 19.**
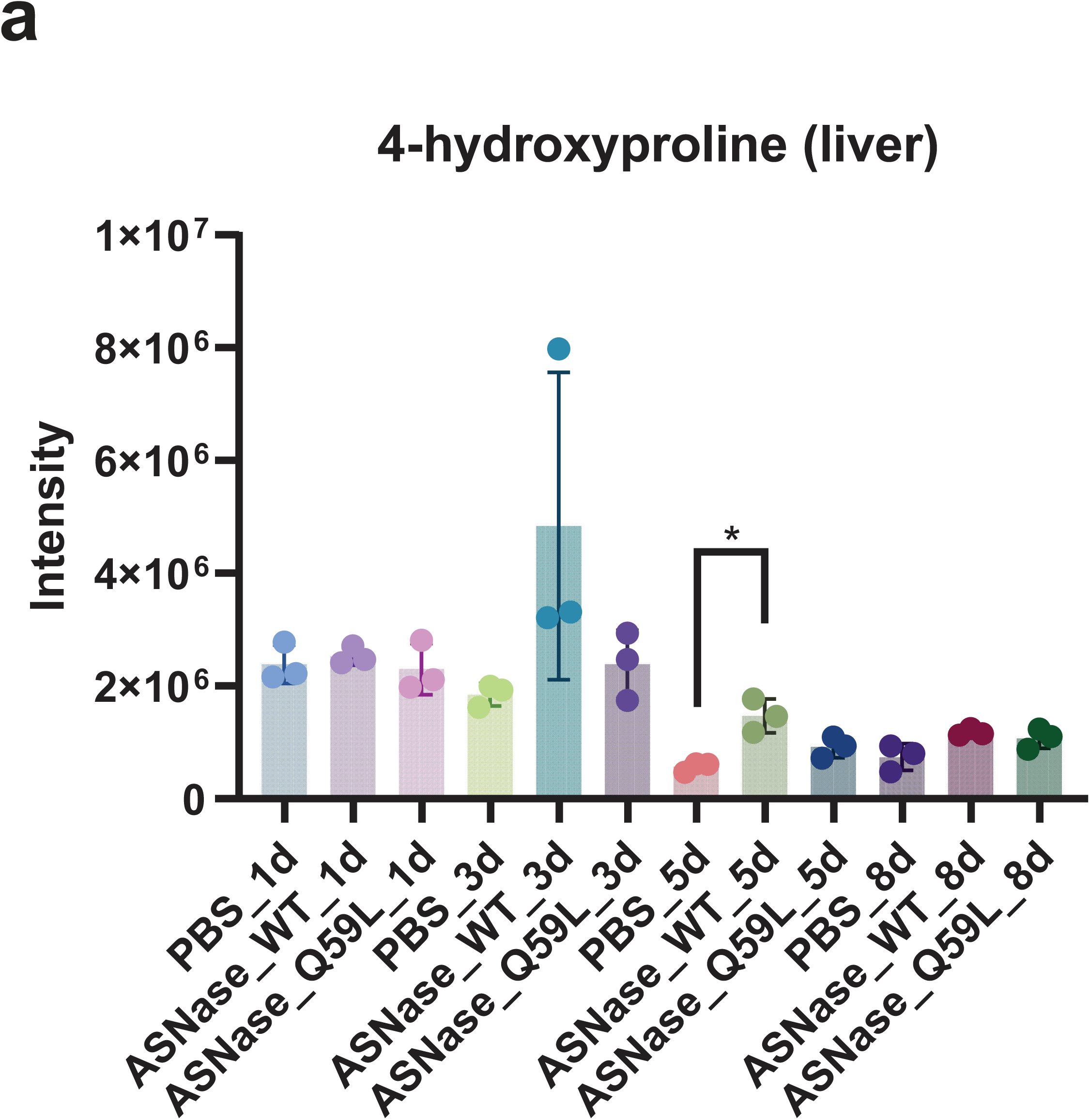
a, The MS intensity of 4-hydroxyproline in the liver under different treatment conditions.

**Supplementary Table 1**, Protein groups and peptides identified using Evosep-Astral and Neo-EV1107-Astral methods.

**Supplementary Table 2**, Protein groups and peptides identified using different MS2 maximum injection time (IT) from 1 to 3.5 ms.

**Supplementary Table 3**, Protein groups and peptides identified using different mass ranges.

**Supplementary Table 4**, Protein groups and peptides identified using different isolation windows from 0.8 to 2 Da.

**Supplementary Table 5**, Protein groups and peptides identified using different AGC values from 100 to 600.

**Supplementary Table 6**, Evaluation of the ultra-fast nDIA method with one to five minutes LC gradients.

**Supplementary Table 7**, Evaluation of the quantitative performance of the ultra-fast nDIA method with three-species samples mixed in various ratios.

**Supplementary Table 8**, Comparison of the protein and peptide identifications using different digestion durations from 30 min to 15 h. To check the missed cleavage percentage, up to three missed cleavage sites were allowed for this analysis.

**Supplementary Table 9**, Protein groups and peptides identified using the standard and high-throughput sample preparation workflows.

**Supplementary Table 10**, Proteins identified in the large-scale mouse tissue proteomics datasets including 507 tissue samples. P-values were calculated using student’s t-test.

**Supplementary Table 11,** Differentially regulated proteins in each tissue following treatment of ASNase for 1 to 8 days. For each protein, the fold change of the median intensity from ASNase treated and PBS treated sample groups was calculated. Only proteins with a fold change >2 or <0.5 and p-value <0.05 were selected as differential proteins.

**Supplementary Table 12,** Number of shared tissues for differential proteins. The differential proteins identified in different time points (1 day, 3 days, 5 days, and 8 days) were merged for each tissue type. The number of shared tissues means certain proteins were identified to be differentially regulated in a certain number of tissues.

**Supplementary Table 13,** Functional clustering analysis of the differentially regulated proteins in the SQWAT according to the online database DAVID (https://david.ncifcrf.gov/).

**Supplementary Table 14,** Functional clustering analysis of the differentially regulated proteins in the GAT.

**Supplementary Table 15,** Functional clustering analysis of the differentially regulated proteins in the lung.

**Supplementary Table 16,** Functional clustering analysis of the differentially regulated proteins in the spleen.

**Supplementary Table 17,** Functional clustering analysis of the differentially regulated proteins in the stomach.

**Supplementary Table 18,** Functional clustering analysis of the differentially regulated proteins in the small intestine.

**Supplementary Table 19,** Functional clustering analysis of the differentially regulated proteins in the large intestine.

**Supplementary Table 20,** Functional clustering analysis of the differentially regulated proteins in the liver.

**Supplementary Table 21,** Functional clustering analysis of the differentially regulated proteins in the kidney.

**Supplementary Table 22,** Functional clustering analysis of the differentially regulated proteins in the heart.

**Supplementary Table 23,** Functional clustering analysis of the differentially regulated proteins in the muscle.

**Supplementary Table 24,** Functional clustering analysis of the differentially regulated proteins in the brain.

**Supplementary Table 25,** Functional clustering analysis of the differentially regulated proteins in the bone marrow.

**Supplementary Table 26,** Metabolomics analysis of amino acid in the tissues.

**Supplementary Table 27,** Metabolomics analysis of polar metabolites in the tissues.

